# Brain states recur during longitudinal viewing and relate to content across narratives and sensory formats

**DOI:** 10.64898/2026.05.31.729141

**Authors:** Yibei Chen, Matin Ghavami, Marie St-Laurent, Lune Bellec, Satrajit S. Ghosh

**Affiliations:** McGovern Institute for Brain Research, MIT, Cambridge, MA, USA; Department of Electrical Engineering and Computer Science, MIT, Cambridge, MA, USA; Centre de recherche de l’Institut universitaire de gériatrie de Montréal, Montréal, Canada; Département de psychologie, Université de Montréal, Montréal, Canada

**Keywords:** brain state dynamics, hidden Markov models, naturalistic paradigm, longitudinal fMRI, functional networks

## Abstract

Models of brain dynamics propose that brain activity revisits a repertoire of configurations, and that current input influences which configuration is expressed. How broadly this holds across a person’s extended experience is unknown. We analyzed 38–57 hours of fMRI per participant in six adults watching the television series *Friends*. Participant-specific hidden Markov models identified 42–47 active brain states, and independent fits to separate portions of viewing recovered similar states with similar tendencies to recur. States matched across participants’ separately fitted models tended to occur at the same moments of the episodes. Among states retained after temporal screening, features from pretrained video, audio, and language networks partly predicted state identity, most strongly from intermediate-to-late layers. Predictions learned from *Friends* generalized without retraining to new films, and language-based predictions extended to reading and audiobook listening. State usage corresponded across stimuli little beyond what each recording’s spectra and covariance reproduced.

## Intro

Everyday experience unfolds through changing sights and sounds, while activity across multiple brain regions varies in relation to this ongoing content (Hasson et al., 2010). Relationships among regional activity time courses also show substantial similarities across rest and different tasks, alongside differences associated with particular tasks (Biswal et al., 1995; Smith et al., 2009; Cole et al., 2014). These relationships describe how regional activity fluctuates together over time, but do not specify how activity is distributed across regions at particular moments. Understanding neural organization during extended experience therefore requires examining both momentary brain states and their organization over time. Which aspects of these states recur across different moments, and how does their expression relate to ongoing content?

Dynamical-systems models treat brain activity as trajectories through changing neural configurations, so both the configurations and the transitions among them are potential levels of organization (Rabinovich et al., 2008; Deco et al., 2011; Breakspear 2017). Whole-brain connectivity configurations also recurred across time and participants, and in single participants they distinguished which task was being performed within windows as short as 22.5 s (Gonzalez-Castillo et al., 2015). From this perspective, recurrence is informative as it tests whether extended experience involves repeated visits to similar distributed configurations rather than a wholly distinct brain state at every moment. Such recurrence would be consistent with the reuse of neural configurations across different moments or events, but would not by itself establish why recurrence occurs, whether it is computationally advantageous, or whether the brain states correspond to discrete biological states. Here, a brain state describes how activity is distributed across regions; its recurrence implies similarity in that distribution, not exact repetition of every region’s activity. A brain state could recur across moments with different content while occurring more frequently under some content conditions than others. Recurrence and content sensitivity therefore describe complementary properties: how broadly a brain state appears across experience, and how its occurrence relates to what a person encounters.

Several lines of prior work describe related, but distinct, forms of brain state recurrence. Event-specific brain states observed during movie viewing were reinstated when participants subsequently recalled the corresponding events, linking initial perception with later recollection (Chen et al., 2017). Similar brain states have also been reported across distinct events that share a schematic structure, including across audiovisual and spoken presentations (Baldassano et al., 2018). In the posterior medial cortex, a small set of brain states identified during movie viewing appeared across multiple movie stimuli and was reactivated during later recall (Lee et al., 2025). Four brain states, estimated from data pooled across participants, occurred in every run of rest, attention tasks, sitcom episodes and a documentary, with occupancy that differed across these contexts (Song et al., 2023). In resting-state recordings, individuals have shown reproducible temporal profiles of recurring co-activation patterns, including the proportion of time spent in particular patterns (Lee et al., 2024). These findings concern different forms of similarity and recurrence; they do not establish how broadly similar brain states, estimated within one person, recur throughout that person’s extended experience. They motivate asking whether similar brain states, with similar tendencies to recur, can be identified in separate portions of such experience.

Interpreting recurrence requires examining the conditions under which brain states occur. Recurring brain states in recorded brain activity may reflect contributions from ongoing content, other biological processes, repeated recording conditions, and properties of the measurement or model. Examining when brain states occur, how long individual occurrences last, and how their prevalence varies across recordings can help characterize their temporal profiles. A brain state concentrated near recording onsets, for example, has a different temporal profile from one expressed throughout recordings, although either could also include content-related activity. These analyses describe observable regularities without uniquely identifying their causes; brain states with no detected timing association remain candidates for content analysis. The distribution of activity across regions further characterizes what each recurring brain state comprises and provides a basis for examining relationships among brain states.

If recurring brain states constitute a repertoire, their organization includes which brain states occur and how activity moves among them. Two experiences could contain the same brain states equally often yet differ in the order in which those brain states occur. Transition structure therefore captures a dimension of temporal organization that cannot be recovered from brain state prevalence alone. Transitions might preferentially connect brain states with similar neural properties, such as their regional activity distributions or their functional-connectivity profiles—the relationships among regional activity fluctuations over time. In resting-state recordings, transition probability was more strongly associated with similarity in functional connectivity than with similarity in regional activation patterns (Vidaurre et al., 2017). This finding motivates asking whether the succession of recurring brain states during extended viewing relates to their similarity, including similarity in how regional activity varies together.

Relating recurring brain states to experience requires distinguishing how often a brain state occurs from when it occurs in relation to content. Comparing overall use across experiences asks whether brain states frequently expressed during one experience also tend to be frequent during another. A separate question is whether stimulus content helps predict which brain state is expressed at a particular moment. Prior work on spoken narratives has linked state expression to acoustic and semantic properties (Liu et al., 2025). During the first viewing of temporally scrambled movie clips, an integrated default-mode and visual state occurred more often at moments that an independent group found comprehensible (Song et al., 2021). Testing a content–state relationship on independent material asks whether it extends beyond the specific experience used to estimate it; connectivity-based models of moment-to-moment emotional arousal, for example, generalized to other movies and other viewers (Ke et al., 2025). Such generalization need not mean that brain states have identical overall prevalence or a fixed one-to-one mapping to content. For example, a brain state associated with speech could occur less often during an experience containing less speech, even if the relationship between speech and brain state expression remained similar. Overall usage and content-based prediction therefore address distinct aspects of how recurring brain states relate to experience.

We asked five related questions. (1) Do brain states recur across separate portions of a person’s extended experience? (2) Do recurring brain states differ in their temporal profiles and regional composition, and do states estimated separately in different people occur at the same moments? (3) How are these states organized in succession, and are transitions related to similarity in regional activity or functional connectivity? (4) Do visual, auditory, and language features predict momentary brain state expression? (5) To what extent do relative brain state usage and content-based prediction generalize to independent experiences? If brain states form a repertoire used in similar proportions whatever the content, relative brain state usage should correspond across experiences while content-based prediction should not generalize. If brain state expression depends on content, content-based prediction should generalize while relative usage need not correspond.

Addressing these questions requires repeated observations within a person across varied experiences. Extended recordings provide opportunities to estimate brain states from separate portions of the data, examine the conditions under which they recur, and characterize their succession. Relating momentary brain state expression to content also requires descriptions of what the person encounters over time. Naturalistic fMRI during audiovisual narratives provides a setting in which brain activity can be aligned with unfolding stimulus features (Hasson et al., 2010). Sampling across episodes and recording sessions allows us to examine whether brain states occur broadly or concentrate in particular portions of experience. A television series nevertheless contains repeated sensory and narrative features, so extensive sampling within it cannot establish how far observed relationships extend beyond that setting. Independent experiences excluded from model fitting are therefore needed to test whether relative brain state use corresponds across experiences and whether content-based predictions apply to new material.

We analyzed longitudinal fMRI recordings acquired while participants watched four to six seasons of *Friends*, fitting a separate hidden Markov model to each participant. The model represents activity across brain regions as a sequence of recurring brain states, which we refer to as states. Models fitted to separate portions of each participant’s viewing recovered similar states with similar tendencies to recur. States differed in their temporal occurrence profiles and regional composition, and transitions preferentially linked states with similar functional connectivity. States matched across participants’ separately fitted models tended to occur at the same moments of the episodes. Among states retained for content analysis, momentary state identity was partly predictable from visual, auditory, and language features of the stimulus, most strongly from intermediate-to-late layers of the feature models. These content–state relationships generalized to varying degrees to independent experiences, including reading and listening for language features, whereas relative state usage showed limited correspondence across experiences. Together, the findings characterize a model-defined repertoire of recurring brain states whose momentary expression relates to ongoing content, while showing that state usage and content-based prediction have distinct limits on generalization.

## Results

### Recurring brain states form a reproducible recurrence gradient across independent subsets of the data

We first assessed whether brain activity while viewing *Friends* repeatedly returned to the same states across the show’s many different scenes, and how widely states varied in their recurring patterns. Six adults each watched up to six seasons of *Friends* while undergoing fMRI (about 146 episodes, split into roughly 292 runs of about 12 minutes each per participant; one participant watched four seasons; **Figure 1A**). For each participant, we processed the fMRI BOLD signals per run and parcellated data from every time point, sampled every 1.49 s, into 156 brain parcels (a combined 100 cortical parcels and 56 subcortical parcels). We reduced the parcellated signal with principal component analysis to the components that captured 95% of its variance (67–77 components), to which we fitted a single hidden Markov model that treated the recording as a sequence of recurring brain states that switch over time (**Figure 1B**). We set a ceiling of 50 states; the model’s prior left states the data did not need unused (**Methods**), and each participant’s data used 37–42 states for more than 1% of all time points.

**Figure 1.**
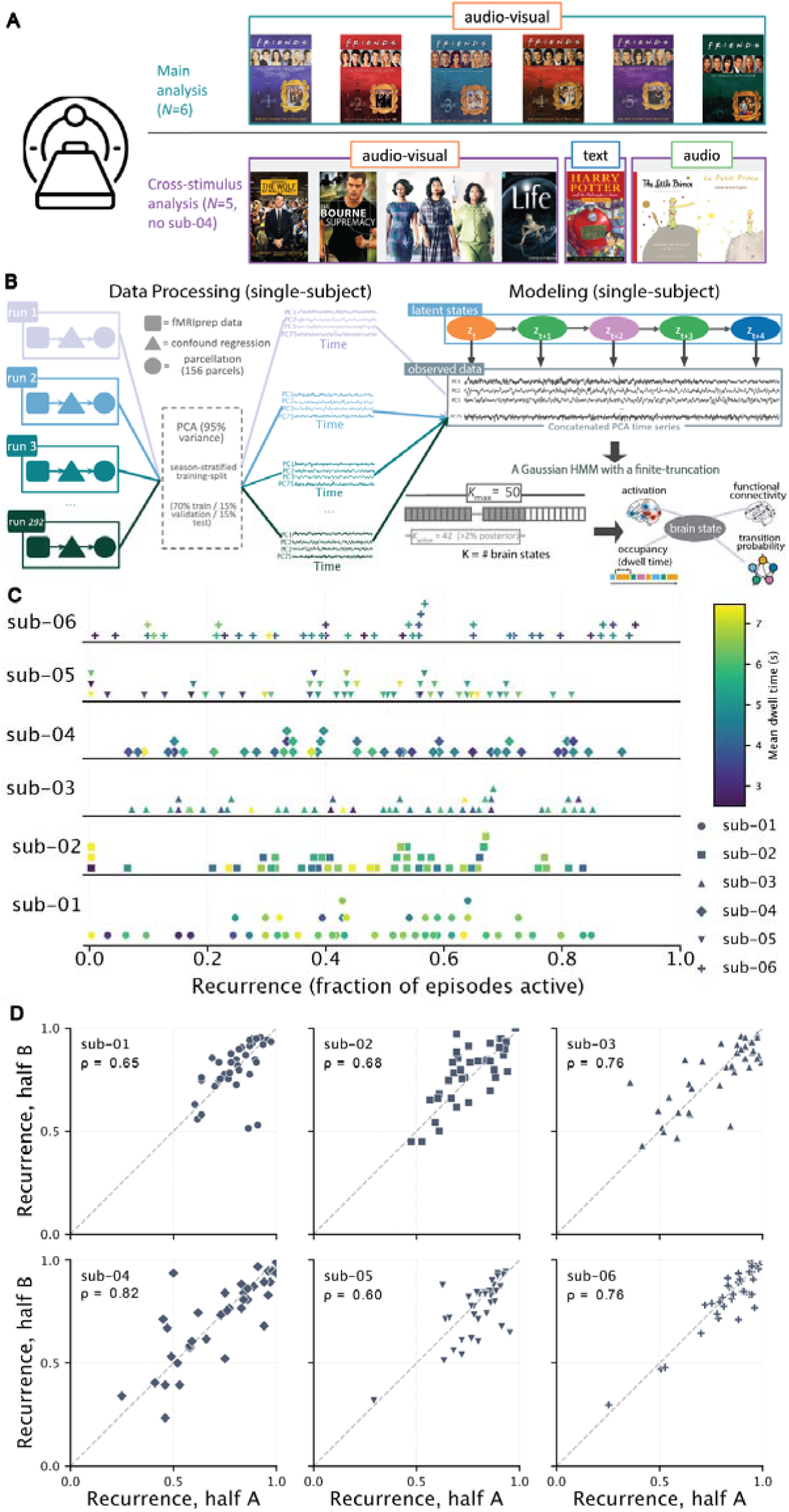
Brain-state recurrence within *Friends* and its reproducibility across independent model fits. (**A**) Datasets. Six healthy adults from the Courtois NeuroMod project watched the television show *Friends* during repeated MRI sessions; sub-04 watched only the first four seasons while everyone else watched six. Most episodes were divided into two ∼12-min runs (a, b) for comfort and to limit head motion, and double episodes into up to four segments (a–d), giving about 146 episodes (∼292 runs) per participant across the six seasons. The same participants also watched ∼10h of films from the CNeuroMod Movie10 set in ∼10-min runs: three feature films (*The Bourne Supremacy*, 2004; *The Wolf of Wall Street*, 2013; *Hidden Figures*, 2016) and one BBC nature documentary (*Life*, 2009); *Hidden Figures* and *Life* were each viewed twice, for 61 runs per participant (*Bourne* 10, *Wolf* 17, *Life* 2×5, *Hidden* Figures 2×12). All participants except subject 4 also read chapter 9 of *Harry Potter and the Sorcerer’s Stone* (1997) in the scanner, and listened to an audiobook of *Le Petit Prince* narrated in English and French (18 runs, 9 per language, ∼9–13 min each; 16 for sub-06). *Friends* is the main analysis (*N* = 6); the other stimuli support the cross-stimulus analysis (N = 6 for *Movie10*; N = 5 for *Harry Potter* and *Le Petit Prince*, no sub-04). (**B**) Processing and modeling. Starting from fMRIPrep derivatives released by CNeuroMod (for *Friends*, fMRIPrep 20.2.3, 20.2.5, or 20.2.6, depending on the sessions, **Table S7**), we applied a minimal confound-regression strategy. Each grayordinate’s time series was z-scored within run before parcellation, so every grayordinate contributes equally to its parcel’s average. Time series were then averaged into a 156-parcel atlas (100 cortical parcels from the Schaefer 100 Yeo 7-network parcellation plus a 56-parcel subcortical and cerebellar set). Per participant, principal-components analysis on the training split (70% train / 15% validation / 15% test, stratified by season) reduced the parcels to latent components capturing 95% of the variance (67–77 components). We then fit one hidden Markov model per participant, treating the PCA time series as generated by a sequence of hidden brain states that switch over time. The model is a Gaussian HMM with a fixed ceiling of *K_max_* = 50 possible states; states the data do not need remain unused. Each used state yields a spatial activation map (its average pattern across parcels), a within-state functional-connectivity pattern (how parcels co-vary while the state is active), how often and how long it is occupied (occupancy and dwell time), and its probabilities of switching to other states. (**C**) Recurrence of states within *Friends*. Each row is one participant; each point is one active state from that participant’s model (sub-01: 46 states, sub-02: 46 states, sub-03: 44 states, sub-04: 44 states, sub-05: 47 states, sub-06: 42 states). The x-axis is the states’ recurrence score: the fraction of *Friends* runs in which the state was active, meaning that the proportion of time spent in this state in a given run (occupancy) exceeded the activity threshold (>2% run duration). Across participants, recurrence scores span from near zero to about 0.9. Points are stacked vertically within narrow recurrence bins to show local density, marked by participant, and colored by the mean duration spent in the state (mean dwell time in seconds). (**D**) Reproducibility across independent fits. For each participant, the *Friends* episodes were split into two interleaved halves, and a separate model was fit to each half. States from the two halves were paired by the similarity of their parcel-space activation maps, and each point gives one paired state’s recurrence score in half A versus half B. The dashed line marks equal recurrence in both halves; ρ is the within-participant Spearman correlation across paired states.

Recurrence, the fraction of runs in which a state is active, is a run-level measure of a state’s prevalence. We calculated a state’s *recurrence score* as the proportion of *Friends* runs in which it occurred. A state was considered present in a run if it occupied more than 2% of the total duration (approximately 14 s). States never present above this level in any run (3–8 per participant) were set aside, leaving each participant an active repertoire of 42–47 states (269 across the six participants), which includes states that exceeded 2% of a run while staying below 1% of the data overall. Within the active repertoire, recurrence scores varied continuously across a broad range, from 0.003 to 0.925; in every participant, the middle 80% of states fell between 0.11 and 0.87 **(Figure 1C**; surface maps of the five highest-recurrence states per participant are shown in **Figure S1**). Some states recurred unevenly across seasons, appearing often in some seasons and rarely in others: a minority in four participants (sub-01: 12, sub-02: 11, sub-03: 8, sub-04: 3) and half or more of the active repertoire in two (sub-05: 34 of 47; sub-06: 21 of 42). Their concentration in particular seasons exceeded what shuffling season labels across runs would produce (5,000 permutations, BH-FDR corrected across the 50 model states; *q* < 0.05).

These recurrence scores were reproducible when the model was refitted on independent portions of the data. Splitting each participant’s episodes into two interleaved halves and fitting a separate model to each, the recurrence scores of matched states (paired between the two fits by the similarity of their region-level activity maps) agreed across halves in every participant (Spearman’s ρ = 0.48–0.84, all *p* < 0.002; **Figure 1D**). Refitting after holding out each season in turn recovered the same state maps in nearly every case (94%–99.6% of matched state pairs per participant had a spatial correlation ≥ 0.3; median spatial correlation of matched states 0.95–0.99; **Figure S4**). The median is far above a derangement baseline, in which each refit state is paired with a state other than its match (null median −0.03, 95th percentile ≤ 0.02), and above each refit state’s best unmatched counterpart (median 0.62–0.72). In 10–19% of matched pairs per participant, the one-to-one assignment did not pair a refit state with its most similar state in the model fitted to all seasons; these pairs form a low tail, so mean similarity (0.87–0.92) is below the median. Because each refit shares all but one season with that model, we treat the match as stability under removal of a season, not as independent replication. Independent component analysis, which has no temporal model, recovered maps close to the states’ maps, slightly better than random rotations of the same principal-component subspace, but did not reproduce the states one by one (**Figure S10**).

### Recurring states differ in the source of their recurrence, and content-eligible states span several brain networks

Apparent recurrence can also arise from measurement artifacts such as head motion and acquisition transients, from the scanning paradigm itself, whose run onsets and slow drift repeat on their own schedule, and from neural variation unrelated to the stimulus, such as changes in arousal (Allen et al., 2014). How much apparent state structure in fMRI reflects neural dynamics at all remains debated (Laumann et al., 2017; Lurie et al., 2020). Therefore, we asked what kinds of states made up each participant’s recurring repertoire by screening every state in the active repertoire and sorting them into different categories (**Figure 2A**; one example per category in **Figure 2D**).

**Figure 2.**
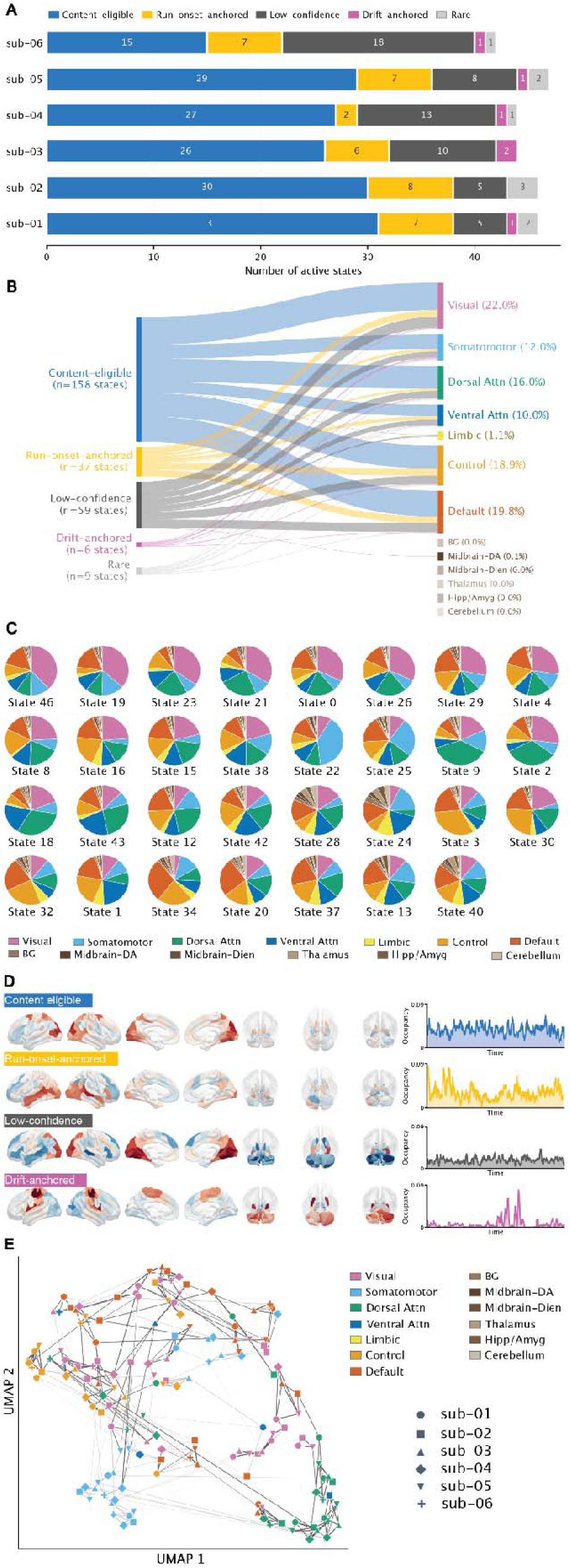
Recurrence categories and brain-network composition of recurring states. (**A**) Per-participant state counts per category over each participant’s active repertoire (42–47 states; rare, low-confidence, run-onset-anchored, drift-anchored, content-eligible; counts inside each segment; x-axis is the number of states), colored as in the key above. Unused states (never above 2% occupancy in any run; 4, 4, 6, 6, 3 and 8 for sub-01 to sub-06) are not shown. (**B**) How each category’s states draw on brain networks. For every state we measured how its activity map spread across 13 brain networks (7 cortical networks from Yeo 7-network plus 6 subcortical groups, including the cerebellum): we took the magnitude (absolute value) of the state’s mean activity in each of the 156 regions, averaged it within each network, and scaled the 13 values to sum to one, giving the state’s network composition. We then kept each state’s three largest networks (rescaled to sum to one) and added these across all states in a category, pooling the six participants. Left nodes are the five categories (with their total state counts); right nodes are the 13 networks; ribbon width is the summed top-three contribution from a category to a network, and each network’s percentage is its share of that total. The pooled ribbons (n = 269 active states) summarize participant-specific states descriptively. Subcortical groups rarely rank among a state’s three largest networks, so their shares are near zero; this reflects the top-three selection rather than an absence of subcortical activity (full 13-network profiles in **Figure S11**). Subcortical node heights are drawn at a minimum size and are not to scale. (**C**) Network composition of sub-01’s content-eligible states. Each pie is one of sub-01’s 31 content-eligible states, showing how that state’s activity map divides among the 13 networks (colors as in the legend below). Pies are ordered by each state’s strongest network (in the legend’s network order) and then by how dominant that network is; the state numbers are the model’s internal labels and carry no meaningful order. Every pie mixes several networks rather than being a single color, the per-state counterpart to the pooled summary in B. (**D**) Example states and their timing, sub-01. One example state per category (content-eligible, run-onset-anchored, low-confidence, drift-anchored; the rare category is not shown). For each example, the brain maps show the state’s mean activity across cortical and subcortical regions, with separate shared color scales for cortical and subcortical maps because their magnitudes differ. The line plot on the right shows the state’s occupancy across sub-01’s full set of runs (the fraction of time spent in the state, binned over the concatenated runs and lightly smoothed); the four strips share one occupancy scale, so categories differ in how their time-in-state is spread out, not in peak height. (**E**) Content-eligible states of all six participants (n = 158), embedded together with UMAP from their activity maps (each relative to the participant’s mean and z-scored across regions; correlation distance). Color shows each state’s strongest network (colors as in C); marker shape shows the participant (as in Figure 1C). Lines join states matched one-to-one between two participants by map correlation (348 matches over the 15 pairs); line width and opacity increase with the correlation. Dark lines mark matches whose two states also occur at the same moments of the episodes both participants watched (228 of 348; **Methods**). Distances in the embedding are not meaningful, and individual lines are descriptive (pair-level tests in **Table S4**; the same matches after removing components shared across participants, all active states, and the embedding under other settings in **Figure S14**).

Categories were assigned in a fixed order of priority (see **Methods**); percentages are of the 269 active states pooled across participants. States that switched too fast for fMRI to resolve, with a typical visit lasting shorter than 3 TRs (about 4.5 s, at or below the limit set by the slow blood-oxygen signal), were labeled *low-confidence* (9.8 per participant, range 5–18; 22%). States whose visits clustered in the first fifth of runs, beyond what shuffling the order of visits within each run would produce (BH q < 0.10), were labeled *run-onset-anchored* (6.2 per participant, range 2–8; 14%). Of the 37 run-onset-anchored states, 19 were anchored at the start of both halves of an episode. Only first halves open with the series theme, so the anchoring of these 19 states cannot be attributed to the theme. Another 15 were anchored only at the start of first halves, where scan onset and the theme coincide, and 3 only at the start of second halves. States whose occupancy (the proportion of time spent in the state in a given run) drifted slowly over the experiment, co-varying with both the season watched and the overall position in the scanning sessions, were labeled *drift-anchored* (1 per participant, range 0–2; 2%). States with none of these flags that were active in fewer than 10% of runs were labeled *rare* (1.5 per participant, range 0–3; 3%). The remaining states, carrying none of these flags and active in at least 10% of runs, were considered *content-eligible* (26.3 per participant, range 15–31; 59% of the active repertoire, 36–67% per participant), and were carried into the network and content analyses.

We projected the recurring states’ activity onto 13 brain networks (7 cortical networks from Yeo 7-network plus 6 subcortical groups, including the cerebellum). Across all five state categories (content-eligible, run-onset-anchored, drift-anchored, low-confidence and rare), states drew mainly on cortical networks (visual, default-mode, control, dorsal- and ventral-attention, and somatomotor), with the limbic network and the subcortical groups contributing little (**Figure 2B**; per-category profiles for all 13 networks are shown in **Figure S11**). Focusing on the content-eligible states, we pooled 158 states identified across the six participants and labeled each one by its most prominent network: the visual network was most commonly identified (44 of 158, 28%), followed by the default-mode (31, 20%), dorsal-attention (29, 18%), somatomotor (27, 17%), and control (22, 14%) networks; only 5 states (3%) were strongly represented in the ventral-attention/salience network, and none in the limbic network.

Few states, however, were dominated by a single network. Across the 158 content-eligible states, the most dominant network accounted for a median of only 25% of a state’s composition (interquartile range 21%–30%), the top three networks for 60% (54%–63%), a median of 4 networks each contributed at least 10% (4–5). The network-spread index, 0 indicating all weight in one network, and 1 indicating weights spread evenly across networks, had a median of 81% across these states (interquartile range 78%–84%), and ranged 79%–83% across participants (**Table S1**). The most common three-network mixtures were visual with control and default-mode networks (31 of 158 states), visual with dorsal-attention and default-mode networks (20), and visual with dorsal-attention and control networks (17). Content-eligible recurring states were therefore not confined to single networks; most states engaged several, spanning visual, attention, default-mode, control, and somatomotor systems (**Figure 2C**).

We further matched content-eligible states one-to-one between participants by their activity maps (**Methods**). Matched states were more similar than chance in all 15 pairs (median *r* 0.60–0.82 against null means of 0.52–0.61; BH-corrected *p* ≤ 0.005) and remained so after each map’s network averages were removed (*p* ≤ 0.009; **Table S4**). Matched states also tended to occur at the same moments of the episodes. Across the episodes both participants watched, the two states’ moment-to-moment occupancy was correlated more than when one participant’s time course was shifted in time (every pair, *p* = 0.001). The correlation peaked at zero lag and fell to zero within about 15 s (**Figure S14E**), and 228 of the 348 matches passed the test individually (dark lines in **Figure 2E**; line lengths in the embedding are not meaningful). Comparable states appearing across participants at the same moments of the stimulus motivate relating them to stimulus content (**Figure 5**).

### Transitions group states into modules and link states with similar functional connectivity

We next asked whether a participant’s states followed one another in an organized way. We first tested whether transitions grouped states into transition modules (sets of states that transition among themselves more often than to states outside the set). We then tested whether transitions linked states with similar functional connectivity, network membership and recurrence.

We measured how strongly each participant’s transitions grouped states with directed modularity Q, the excess of transitions within groups over what each state’s overall rates of leaving and entering predict (Leicht & Newman, 2008). We found the grouping that maximized Q with the Louvain algorithm (Blondel et al., 2008). Transitions counted from the decoded state sequences were compared with two kinds of sequences that keep a simpler structure. In the first, the order of visits was shuffled within each run, which keeps which states occurred in each run and how long every visit lasted. In the second, sequences were simulated from a chain without modules that keeps each state’s tendency to stay and its popularity as a destination. In all six participants, Q was 0.36–0.47, exceeding the mean of the shuffled-order null by 0.21–0.38 and that of the chain null by 0.26–0.41 (p = 0.001 and 0.002, the floors for 1,000 and 500 draws).

Modular structure was present in held-out seasons, but module membership was reproduced only in part. Transitions in each held-out season, decoded by a model fitted without that season, were more modular than their shuffled-order nulls in all 34 folds (excess 0.17–0.38; *p* = 0.001 in every fold). The partition of those held-out transitions agreed with the main partition above chance in every fold (adjusted Rand index 0.22–0.85, where 0 is chance agreement and 1 is identical partitions; participant medians 0.35–0.62; permutation *p* ≤ 0.0008), but well below identity. Within the full data, the partitions were stable across bootstrap resamples of runs (median adjusted Rand index 0.61–1.00). The model’s fitted transition matrix is divided into 4–6 modules per participant (**Figure 3A**). These partitions agreed with those of the counted transitions (adjusted Rand index 0.74–1.00 in five participants and 0.58 in sub-05).

**Figure 3.**
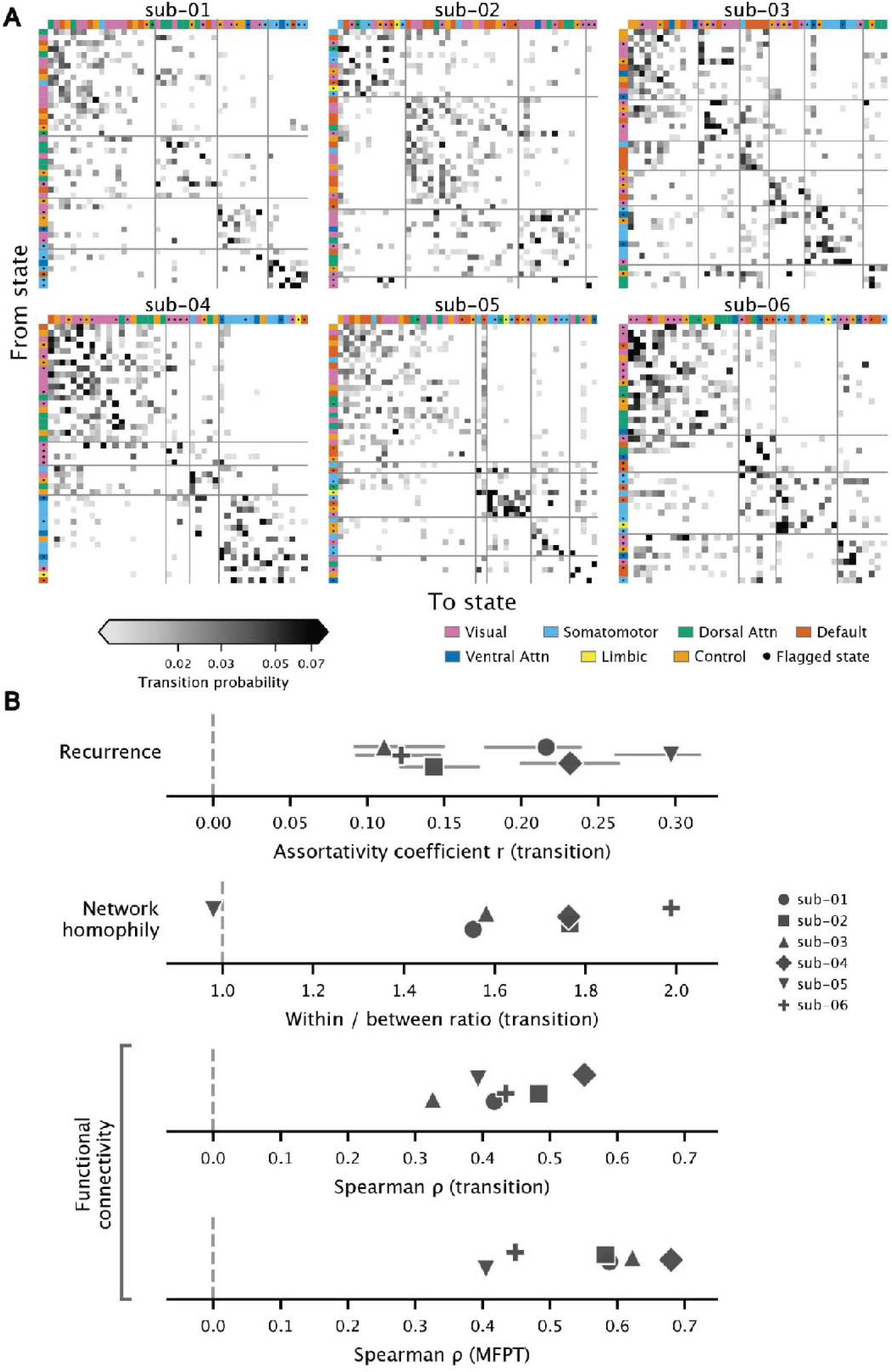
State transitions vary with recurrence, network membership, and functional-connectivity similarity. (**A**) Per-participant state transition matrices, ordered by transition module. Each cell is the fitted probability of moving from the state in its row to the state in its column (the model’s transition matrix, self-transitions removed); darker means a more likely transition, on a logarithmic grey scale between the 5th and 95th percentiles of the displayed probabilities across participants (0.010–0.074; stronger cells saturate), with probabilities below 0.01 left white. States are ordered along both axes by the partition of this matrix with the highest modularity: modules are sorted by the recurrence of their most recurrent state, and states within a module by recurrence. Mid-grey lines separate modules. Colored ribbons on the top and left edges give each state’s dominant network (the same colors as Figure 2), and dots on the ribbons mark states excluded from content analyses. Functional-connectivity and activation-map similarity in the same state order are shown in Figure S15. Modularity was tested on the transitions counted from the decoded sequences, whose partitions agree with those shown (adjusted Rand index 0.74–1.00 in five participants, 0.58 in sub-05). Q of the counted transitions, with its excess over the mean of the shuffled-visit-order null (1,000 draws) and of the non-modular-chain null (500 draws): sub-01 0.44 (+0.24, +0.31), sub-02 0.43 (+0.21, +0.26), sub-03 0.36 (+0.27, +0.28), sub-04 0.39 (+0.26, +0.28), sub-05 0.47 (+0.29, +0.34), sub-06 0.45 (+0.38, +0.41); *p* = 0.001 and 0.002 in every participant, the floors for those draw counts. (**B**) Four measures of how the transitions are organized, each on its own axis, one marker per participant (n = 6). Recurrence assortativity measures whether states with similar recurrence scores tend to transition into one another (Newman (2003)’s assortativity coefficient, positive when similar-recurrence states are preferentially linked; grey whiskers are a bootstrap 95% interval; sub-01=0.216, sub-02=0.143, sub-03=0.111, sub-04=0.232, sub-05=0.297, sub-06=0.122). Recurrence closely follows each state’s long-run occupancy under the fitted transition matrix (Spearman’s ρ = 0.97–0.98 with its stationary distribution), so assortativity describes how transitions concentrate on frequently occupied states rather than relating two independent measures. Network homophily is the average transition probability, over state pairs with at least one observed transition, between states that share a dominant network divided by the same average between states with different dominant networks (1 means transitions ignore network membership; above 1 means within-network transitions are more likely; sub-01=1.55, sub-02=1.77, sub-03=1.58, sub-04=1.76, sub-06=1.99). The functional-connectivity measure asks whether state pairs that transition between each other more often also have more similar within-state functional connectivity, where a state’s functional connectivity is the pattern of co-activation among brain regions while that state is active and the similarity of two such patterns is summarized by the RV coefficient (a correlation-like index from 0 to 1, with 1 meaning identical patterns); the association is a Spearman correlation across state pairs (sub-01=0.417, sub-02=0.483, sub-03=0.326, sub-04=0.551, sub-05=0.393, sub-06=0.437). The mean-first-passage-time measure asks whether states that take more transition steps to reach from one another also have less similar connectivity, where mean first-passage time is the expected number of steps to travel from one state to another (from the model’s fitted transition matrix); again a Spearman correlation (sub-01=0.588, sub-02=0.582, sub-03=0.622, sub-04=0.680, sub-05=0.405, sub-06=0.431). Vertical dashed lines mark each measure’s null (0 for the correlations and assortativity, 1 for the ratio). Assortativity (0.11–0.30) and the FC (0.33–0.55) and MFPT (0.40–0.68) correlations were positive in every participant (permutation *p* ≤ 0.0012); network homophily exceeded 1 in five participants (1.55–1.99, *p* ≤ 0.0084) and not in sub-05 (downward triangle; 0.98, *p* = 0.51).

Activity stayed within a module for longer than it would have if the same state visits had come in random order. Consecutive time in one module averaged 5.5–8.9 TRs (8.1–13.2 s), against 4.1–5.9 TRs when the order of visits within each run was shuffled (p = 0.001 in every participant, the floor for 1,000 draws); a single state visit averaged 2.6–3.8 TRs. Modules also tended to be occupied at the same moments across participants. We matched each participant’s modules to another participant’s by their mean activity maps, without using their timing. For each matched pair, we then correlated, TR by TR over the runs both participants had watched, whether each participant was in that module. The mean correlation over matched modules was 0.03–0.21 per participant pair, against −0.003 to 0.003 when one participant’s runs were circularly shifted in time (p = 0.001, again the floor, in all 15 pairs).

Modules did not reproduce the canonical networks. Module membership agreed weakly with each state’s dominant network (adjusted Rand index 0.03–0.18), and 25 of the 27 modules contained states with different dominant networks (**Figure 3A**). We next asked whether relations between pairs of states that do not depend on the order of visits could account for the modules. For each participant, we predicted every transition probability between states with a defined functional connectivity (four states without one were excluded) from seven pairwise measures: the similarity of the two states’ functional connectivity, the correlation of their activation maps, the overlap of their activity distributions, whether they shared a dominant network, their difference in recurrence, and their co-occurrence across runs and across seasons (how closely their occupancy rose and fell together). The prediction also allowed for how often each state was left and entered. In no participant was the fitted transition matrix more modular than 1,000 matrices built from these predictions with the prediction errors shuffled across state pairs (*p* = 0.06–0.97).

To see which of these relations the modules followed, we compared pairs of states in the same module with pairs in different modules. Pairs in the same module had, on average, more similar functional connectivity and, in five of six participants, more similar activation maps (**Figure S15**) and higher co-occurrence across runs. When all seven measures were entered together, co-occurrence across runs was the strongest predictor of sharing a module in every participant (p = 0.0002, the floor for 5,000 permutations); activation-map similarity contributed less consistently, and connectivity similarity made no consistent contribution (**Table S5**). We next related these measures to transitions between individual pairs of states.

State pairs that transitioned between each other more often had more similar within-state functional connectivity in all six participants (Spearman’s ρ = 0.33–0.55, *p* = 0.0002, the floor for 5,000 permutations; **Figure 3B**). The association was not explained by connectivity shared across all states, by state size or by activation-map similarity. After subtracting the mean connectivity across states and partialling out the two states’ TR counts and the correlation of their activation maps, it remained positive in every participant (partial Spearman ρ = 0.16–0.42, against 0.33–0.55 for raw RV; permutation p = 0.0002, the floor for 5,000 permutations). States that took more steps to reach from one another (a longer mean first-passage time) were less similar in connectivity, also in all six participants (ρ = 0.40–0.68, *p* ≤ 0.0012). In five participants, transitions were more likely between states sharing a dominant network than between states with different dominant networks (within- to between-network ratio 1.55–1.99, *p* ≤ 0.0084). In sub-05 the ratio was 0.98 (p = 0.51), although its connectivity effects were like those of the other participants. Transitions also linked states with similar recurrence (assortativity 0.11–0.30, *p* ≤ 0.001 in every participant). Recurrence closely followed each state’s long-run occupancy under the fitted transition matrix (Spearman’s ρ = 0.97–0.98 with its stationary distribution). This assortativity therefore describes how transitions concentrate on frequently occupied states, not a relation between two independently measured properties. Per-participant functional-connectivity and network-homophily values are shown with each participant’s audio decoding depth in **Figure S9**.

In every participant, transitions grouped states into modules that were occupied for longer stretches than a random order of visits produces, and at the same moments across participants. The modules cut across the canonical networks and were no more modular than pairwise relations between states predict. Transitions between individual pairs of states followed the similarity of their functional connectivity.

### A phase-randomized null reproduces most of the correspondence between *Friends* recurrence rank and film occupancy, and most reading and listening correlations sit at or below it

We next asked whether the ordering of states by recurrence also held for stimuli other than *Friends*. We applied each participant’s *Friends*-trained pipeline (PCA and HMM) to the other stimuli without retraining and, for each target stimulus, correlated every active state’s mean occupancy (the average fraction of a run spent in that state) with its *Friends* recurrence score (**Figure 4**).

**Figure 4.**
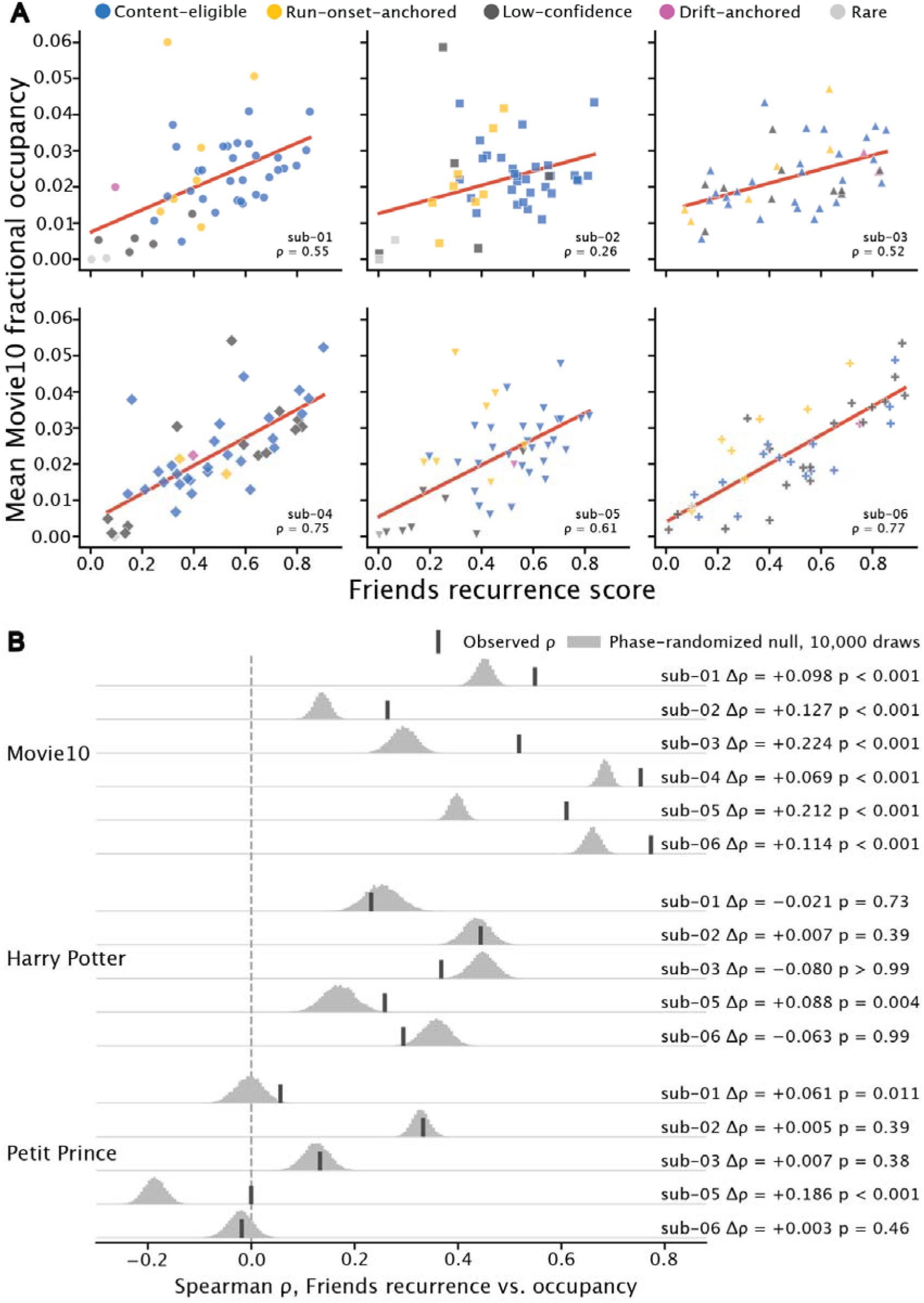
*Friends* recurrence rank versus occupancy under other stimuli. **(A)** Per-participant correspondence to *Movie10*. Each subplot is one participant; each point is one active state from that participant’s *Friends*-trained model, placed by its *Friends* recurrence score (x) and its mean fractional occupancy when the participant’s *Friends*-trained pipeline (PCA and HMM) was applied to *Movie10* without retraining (y). Points are colored by recurrence category (Figure 2) and shaped by participant. The annotated ρ is the within-participant Spearman correlation between recurrence score and *Movie10* occupancy; the red straight-line fit is a visual guide. **(B)** Observed correlation against its phase-randomized null, by stimulus: *Movie10* (audiovisual films, 61 runs per participant), *Harry Potter* (word-by-word visual reading, no sound, 7 runs) and *Le Petit Prince* (audiobook listening, no picture, 16–18 runs, French and English pooled). Each row is one participant; sub-04 contributed no *Harry Potter* or *Le Petit Prince* data. The grey ridge is the distribution of the same Spearman correlation across 10,000 surrogates in which that stimulus’s runs were phase-randomized, drawn as a histogram on shared bins and scaled to a common peak height. The dark tick marks the observed correlation (for *Movie10*, the value annotated in A). Each row is labeled with Δρ, the observed correlation minus the null mean, and the one-sided empirical *p* (minimum 1/10,001, shown as *p* < 0.001). The dashed line marks zero correlation. For *Le Petit Prince* sub-05 the null lies below zero (mean −0.19) and the observed correlation is near zero, so its margin above the null does not reflect a positive correlation.

*Friends* recurrence rank correlated positively with *Movie10* occupancy in all six participants (Spearman’s ρ = 0.26-0.77: sub-01 0.55, sub-02 0.26, sub-03 0.52, sub-04 0.75, sub-05 0.61, sub-06 0.77; **Figure 4A**).

A positive correlation could arise even if the states did not correspond meaningfully across *Friends* and the films. Every stimulus was analyzed using the same *Friends*-trained model and data from the same participant, and recordings from the same brain share statistical properties across stimuli, such as the power spectrum of each principal component and the covariance among components. Applied to signals with these properties, the model may favor some states over others, and that uneven occupancy alone could correlate with recurrence during *Friends*.

To estimate this baseline, we generated 10,000 surrogates of the *Movie10* runs by phase randomization. The surrogates preserved each run’s power spectra and covariance but disrupted its moment-to-moment dynamics, and we recomputed the correlation for each. The surrogates produced substantial correlations: the mean null correlation ranged from ρ = 0.14 to 0.68 across participants, 52–91% of the observed value. The observed correlation nevertheless exceeded its surrogate distribution in every participant, by a modest margin (Δρ = 0.07–0.22; *p* = 0.0001, the minimum possible with 10,000 draws; z = 5.97–15.62; **Figure 4B, Figure S13** and **Table S2**). Δρ measures how far the observed correlation lies above the null mean; Spearman correlations do not divide into additive parts, so Δρ is not a separate, stimulus-specific component.

The correlation between recurrence during *Friends* and occupancy during *Movie10* differed across the four films. Averaged across participants, it was ρ = 0.69 for *The Wolf of Wall Street* (drama), ρ = 0.63 for *Hidden Figures* (drama), ρ = 0.40 for *The Bourne Supremacy* (action), and ρ = 0.16 for *Life* (nature documentary). This ordering of the participants matched the order of the films’ mean log-likelihood gaps, from smallest to largest (**Figure S8B**), although the two orders agreed in only four of six participants. The phase-randomized null pooled all *Movie10* runs, so the films were not tested against it individually, and we do not interpret the ordering by genre.

We also examined two stimuli presented in a single modality: *Harry Potter*, word-by-word visual reading without audio, and *Le Petit Prince*, an audiobook without visual input. Across the five participants with available data (sub-04 had no data for either), recurrence during *Friends* correlated with occupancy during *Harry Potter* at ρ = 0.23–0.44 (mean ρ = 0.32) and during *Le Petit Prince* at ρ = −0.02 to 0.33 (mean ρ = 0.10). The same phase-randomized null, run on each stimulus’s own runs, had means of ρ = 0.17–0.45 for *Harry Potter* and −0.19 to 0.33 for *Le Petit Prince*. For *Harry Potter*, the observed correlation sat above its null in one participant (sub-05, z = +2.77), near it in two (z = −0.64 and +0.29) and below it in two (z = −2.41 and −3.22). For *Le Petit Prince*, it sat above its null in two participants (sub-01, z = +2.24; sub-05, z = +9.08, where the null lay below zero and the observed correlation was near zero) and at it in three (z = 0.11–0.29; **Figure 4B**).

Sub-02 had the weakest *Movie10* correlation (ρ = 0.26) but the largest raw correlations for reading and listening (ρ = 0.44 and 0.33), both close to their null means (z = +0.29); correlations on content-eligible states alone are in **Table S3**.

Applied to resting-state recordings from all six participants, the *Friends*-trained components reconstructed rest about as well as *Friends* (*R²* = 0.94–0.96), but the model fit rest worse than held-out *Friends* in every participant (**Figure S8**). Without a phase-randomized null for rest, we did not interpret its correlations.

Overall, the ordering of states by *Friends* recurrence corresponded to *Movie10* occupancy in every participant, and in every participant the observed correlation exceeded a null that preserved each run’s power spectra and covariance, by a modest margin. For word-by-word reading, the observed correlation sat above its null in one participant, near it in two and below it in two; for audiobook listening, it sat above its null in two participants and at it in three.

### Content-eligible state identity is decodable from video, audio, and language features of the stimulus, with peak depth differing by model and preserved in new films for video and audio

As a final step we asked whether states visited by the brain were related to what each participant was seeing and hearing while movie watching. We trained a classifier to decode which content-eligible state the brain was in at each moment in time, using corresponding stimulus features extracted from frozen pretrained video, audio, and language models. Features were used one layer at a time to locate the model depth at which decoding peaked (**Figure 5**).

**Figure 5.**
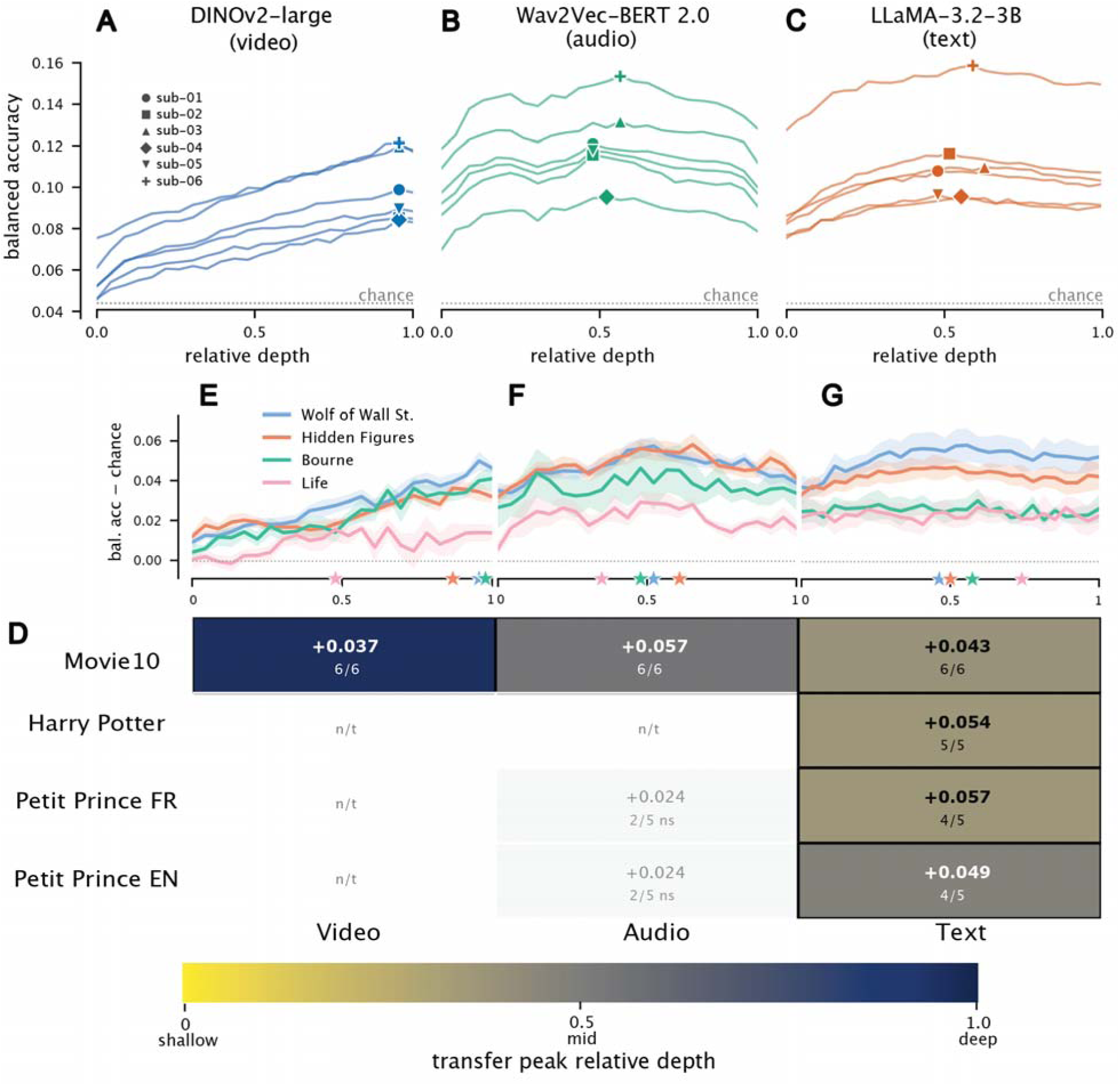
Decoding fitted-state identity from transformer-model features, layer by layer. (**A-C**) Within-*Friends* decoding. For each participant, a classifier was trained to predict which content-eligible state the brain was in at each time point using layer-specific stimulus features from each of the following frozen pretrained models: DINOv2-large for video (**A**, 24 layers), Wav2Vec-BERT 2.0 for audio (**B**, 24 layers), and LLaMA-3.2-3B for text (**C**, 28 layers). Each curve reflects one participant’s decoding accuracy (balanced accuracy, which corrects for states being visited for unequal amounts of time) as a function of model depth, taken at the brain-to-feature lag that decoded best for that participant (lags of 0 to 8 TRs were tested; lag 0 was held out as a synchrony check); markers show each participant’s peak layer (project-wide subject shapes). The dotted line is the mean chance level. Significance used a within-run circular-shift null (shifting each run’s state sequence in time to break the brain-feature correspondence while preserving its temporal structure), corrected across the lag-by-layer grid (Benjamini-Hochberg); a timing-only baseline (six regressors capturing run-onset and session-position effects) served as a decoding floor. The y-axis (absolute balanced accuracy, not accuracy above the floor) is shared across A-C so magnitudes can be compared across modalities. The x-axis is relative model depth: each model’s layers are rescaled to run from 0 (the first, shallowest layer, nearest the input) to 1 (the last, deepest layer, nearest the output), so depth can be compared across models with different layer counts. (**D**) Generalization of the depth pattern to other stimuli. Each *Friends*-trained decoder was applied, without retraining, to the other stimuli the same participants viewed (the *Movie10* films, *Harry Potter*, and *Le Petit Prince* in French and English; see Figure 1A), wherever the matching modality was present. Rows are target stimuli, columns are modalities. Outlined cells, colored by depth (colorbar below), mark combinations where a majority of available participants passed FDR correction at their peak: the color is the median peak relative depth and the number is the median peak balanced accuracy above chance. Muted cells without an outline mark tested combinations where fewer than half passed (a single noisy peak is not shown as a replicated depth). Hatched cells (n/t) mark combinations that could not be tested because the modality was absent, such as video for the audio-only or text-only stimuli. (**E-G**) *Movie10* generalization by film. Each colored line is one *Movie10* film, showing mean decoding accuracy above chance across participants over relative depth, with shaded bands giving the standard error across participants; the star on the depth axis marks each film’s median peak layer. A shared legend names the four films, and the y-axis is shared across E-G.

Within *Friends*, features from all three models predicted state identity above two baselines in every participant. The first baseline captured how accurately states could be predicted from timing alone (**Figure S6**), which is from a moment’s position within the run and session. The second was a shuffled null, created by shifting each run’s state sequence in time. This shift broke the alignment between states and stimulus features while preserving the temporal structure of the state sequence.

Video-based decoding was most accurate near the end of the video model, peaking at layers 22 of 24 in all six participants (relative depth 0.96). Peak balanced accuracy ranged from 0.084 to 0.124, compared with participant-specific chance levels of 0.032–0.067, and the best lag was 3 TRs (approximately 4.5 s; **Figure 5A**). Audio-based decoding peaked near the middle of the audio model, at layers 11–13 of 24 (relative depth 0.48–0.57; peak balanced accuracy 0.096–0.159; best lag 3–4 TRs; **Figure 5B**). Text-based decoding also peaked near the middle layers in all six participants (peak balanced accuracy 0.095–0.164; best lag 3 TRs; **Figure 5C**), and varied less across layers than video decoding: the best layers, typically 13–17 of 28 (relative depth 0.48–0.63), only slightly outperformed the others.

For video, the first transformer layer decoded near the timing floor (normalized effect size 0.006–0.021, against 0.004–0.017 for the timing-only baseline and 0.049–0.087 at the peak layer). The late-layer advantage is therefore not explained by information already available after the first layer. For audio and text the first layer already decoded well above the timing floor (0.033–0.076), and the gain from the first to the peak layer was smaller than for video in every participant (0.021–0.041 against 0.042–0.066).

We next tested whether the late video-model peak was limited to states dominated by the visual network. At each participant’s best lag, we repeated the analysis within groups of states that shared a dominant network and the sign of their loading on it. Twelve such groups across five participants had enough states; none of sub-06’s did. Decoding again peaked in later video-model layers (layers 16–23 of 24) for groups dominated by the default-mode and dorsal-attention networks as well as visual ones, so the late peak was not limited to visually dominated states (**Figure S5**).

We then asked whether the relationship learned from *Friends* generalized to new material. Each participant’s *Friends*-trained brain-state model first assigned a state to every moment of *Movie10*, which consisted of three feature films and a nature documentary, with the same audiovisual format as *Friends* but different casts, settings, and storylines. The same pretrained models extracted video, audio, and language features from *Movie10*. Without retraining, a classifier fitted to all *Friends* time points at each participant’s best *Friends* lag then predicted the states assigned during *Movie10*.

All three decoders predicted state identity in *Movie10* in all six participants after correction. The layer pattern was preserved for video and largely for audio. Video decoding again peaked in later layers (layers 22–23 of 24; relative depth 0.96–1.00), within one layer of each participant’s *Friends* peak. Audio decoding peaked near the middle (layers 11–16 of 24; relative depth 0.48–0.70), within two layers of the *Friends* peak in five participants and five layers later in sub-05. Text decoding peaked earlier than in *Friends* in five of six participants and varied more across participants (layers 7–14 of 28; relative depth 0.26–0.56; **Figure 5D**). Per-layer profiles for every held-out stimulus are shown in **Figure S7**.

Generalization to the other stimuli depended on the type of decoder. We applied the audio decoder to the French and English versions of *Le Petit Prince*. Of the ten participant-by-language combinations (five participants and two languages; no data were available for sub-04), four were significant after correction: sub-02 and sub-03 in both languages. In the other six, decoding did not exceed the shuffled null at any layer after correction. The text decoder generalized more consistently to the language-based stimuli. It predicted state identity during *Harry Potter* in all five participants with available data and during *Le Petit Prince* in eight of the ten participant-by-language combinations, four of five in each language; sub-05 was not significant in either. The layer at which text decoding peaked varied across participants and stimuli (layers 9–13 of 28 for *Harry Potter* and 4–17 in the significant *Le Petit Prince* combinations). State-identity prediction from language features therefore generalized to reading and listening, but not at a consistent layer depth. Video decoding could not be tested for *Harry Potter*, which consisted of word-by-word reading, or for *Le Petit Prince*, which contained audio only.

An exploratory analysis of the four *Movie10* films showed that video decoding peaked later in the model for three feature films (median relative depth 0.96 for *The Wolf of Wall Street*, 0.87 for *Hidden Figures* and 0.98 for *The Bourne Supremacy*) and earlier for the nature documentary *Life* (0.48), where the peak layer was significant in only one participant (**Figure 5E**).

Overall, state identity could be predicted from stimulus features both within *Friends* and, to varying degrees, across new stimuli. Decoding exceeded the shuffled null, and its strongest layer differed by model that late layers for video and middle layers for audio and text. The moment-to-moment relation between stimulus features and state identity extended to new material, whereas the ordering of states by recurrence was largely reproduced by surrogates that kept only each recording’s spectra and covariance.

## Discussion

Across tens of hours of varied viewing, each participant’s brain activity returned to a reproducible repertoire of states, estimated from that person’s data alone. Models fitted to separate portions of viewing recovered similar states with a similar ordering of recurrence. Transitions grouped states into modules and linked states with similar functional connectivity, and matched states in different participants tended to occur at the same moments of the episodes. Which state was expressed at a given moment was partly predictable from the video, audio, and language content being viewed, including in material the models had not been fitted to. How often each state was used corresponded across stimuli little beyond what each recording’s spectra and covariance reproduced.

Broad recurrence does not imply that a state is independent of input. A state can recur across hours of viewing and still be expressed more when particular content is present. Functional organization is stable within individuals (Gordon et al., 2017; Gratton et al., 2018; Lee et al., 2024; Hermosillo et al., 2024; Dworetsky et al., 2024), including during movie viewing (Vanderwal et al., 2017), yet which states are expressed, and how often, changes with the context in which the same story is heard (Chen et al., 2026). Both can hold if occupancy partly reflects a person’s content–state relationship applied to the features of a particular experience. In our data, the content–state relationship estimated during *Friends* predicted state identity in films and, to varying degrees, in reading and listening, whereas the ordering of state use exceeded its null in every participant only for films. The two analyses differed in measure, state set, and participant coverage, so they do not show which relationship is more stable.

This distinction matters for occupancy and dwell time, the usual summaries for comparing state dynamics across sessions and individuals (Vidaurre et al., 2017; Ahrends et al., 2025). If state expression depends partly on content, an occupancy difference between two conditions or groups can arise from the experiences sampled, with no change in how a person’s states relate to content. Two tests can separate these: a decoder trained in one condition should predict the state sequence of another whose occupancy differs, and occupancy differences should shrink when the two experiences’ feature distributions are matched. Raw occupancy similarity can also overstate correspondence. Recurrence closely followed each state’s long-run occupancy under the fitted transition matrix, so the model’s general preference for some states produces a positive correlation by itself. A null preserving only spectra and covariance reproduced half to nine-tenths of the correspondence between *Friends* and *Movie10* and, in most participants, correlations as large as the observed ones for reading and listening.

State identity was predicted best from layers beyond the first, late in the video model and at intermediate depth in the audio and language models. For video, the first layer predicted state identity near the level of timing alone, and the late peak was not limited to states dominated by the visual network (**Figure S5**). For audio and language, the first layer already predicted state identity well, and the gain at the peak was smaller. Encoding studies predict regional responses to natural speech and language from such models (Goldstein et al., 2022; Caucheteux et al., 2023; Antonello et al., 2023), and middle layers of speech models best predict the auditory cortex (Vaidya et al., 2022). Those studies predict regional activity; here the target was the fitted state, so the result concerns which layers carry information about state identity, not which regions represent which features.

The language decoder, trained on dialogue heard while watching, also predicted state identity during word-by-word reading in all five participants with data and during audiobook listening in 8 of 10 participant-by-language combinations, so the relation it learned did not depend on a single sensory format. Nor did it rest on whether speech was present. With periods without words excluded, accuracy stayed above the null in all 21 participant-by-stimulus combinations and was significant in 20 (**Table S11**). The audio decoder predicted state identity in 4 of the 10 audiobook combinations. The audiobook’s single narrator could explain the difference; the number of runs cannot, because the language decoder succeeded on the same runs. A hierarchy of temporal receptive windows, in which regions farther from sensory input integrate over longer stretches of a narrative (Hasson et al., 2008; Lerner et al., 2011), would allow state expression to relate to content beyond single images or sounds. The late video peak and the generalization across formats are compatible with that hierarchy but do not test it: deeper layers retain low-level information, the video features came from single frames, each modality had one model, so depth and architecture are confounded, and layer depth was not related to any cortical timescale.

Brain activity returned to the same patterns across episodes viewed hours to days apart, rather than taking a new pattern at every moment. A finite-state model assigns repeated labels by construction, so the evidence for reuse lies elsewhere: most states were active in many runs, and models fitted to separate halves of a participant’s runs reproduced the ordering of recurrence. Such return differs from event-specific patterns, which hold within an event, change at its boundaries (Zacks et al., 2007; Baldassano et al., 2017; Geerligs et al., 2022; Henderson et al., 2025) and return when the event is recalled (Chen et al., 2017). States here returned in different episodes, and their return does not by itself imply the schematic organization shared by events with a common structure (Baldassano et al., 2018). Recurrence was uneven across seasons for a minority of states in four participants and for half or more in two. Season and preprocessing release were partly confounded, and the test that separates them had low power and could not be run for sub-06 (**Methods**).

About two-fifths of each active repertoire was set aside before the content analyses: states whose occupancy concentrated at run onsets or drifted across sessions, and states whose visits were too short or too sparse to characterize. Screening used timing alone, so the retained states were not selected on the spatial and stimulus relations tested later, and it does not separate neural signals from artifacts. Independent component analysis matched the state maps only slightly better than random rotations of the same subspace and did not reproduce the states one by one (**Figure S10**). The reproducible organization therefore lies in each participant’s subspace and recurrence ordering rather than in a particular decomposition, and no fitted state should be read as a uniquely defined biological entity. A bounded repertoire is compatible with descriptions of population activity as low-dimensional and constrained (Shine et al., 2019; Vyas et al., 2020), but the data do not establish the attractor or energy-landscape structure those descriptions propose (Hopfield 1982).

Separately fitted repertoires were aligned across people. Matched states were more similar than under a null that mixes each repertoire’s states, by a modest margin, and they occurred at the same moments of the episodes, with occupancy correlations that peaked at zero lag. This time-locking is consistent with inter-subject synchronization during natural viewing (Hasson et al., 2004), here between states estimated without pooling data across people. The succession of states was organized as well. Transitions grouped states into modules that held activity longer than a random order of visits, cut across canonical networks, and were no more modular than pairwise relations between states predict (**Figure S15; Table S5**). Modular structure was present in every held-out season, although module membership was reproduced only in part. Transitions also linked states with similar functional connectivity, as in group-level resting-state models (Vidaurre et al., 2017), and the association remained after allowing for connectivity common to all states, state size and activation-map similarity.

Several limits apply. Six participants cannot establish prevalence or how repertoires differ between people; recording 38 to 57 hours per person required that trade, and larger samples will be needed to test between-person variation, for example with age (Lugtmeijer et al., 2025). The repertoires came from one television series. Films, reading, listening and rest extended the comparison, but those conditions were mostly narrative and differed in run count and participant coverage. Recurrence was inferred from a hemodynamic signal, so only states resolvable at that timescale could be detected, and autocorrelation could raise transition probability and connectivity similarity together. An autocorrelation-preserving null and electrophysiological recordings would address this.

Decoding and correspondence were associative: each exceeded a null, but no analysis manipulated the stimulus, and decodability does not show that a state represents particular content. State number, boundaries and maps depend on the model, including its first-order transitions. Refits without each season recovered the same state maps but shared all but one season with the full model, so they show stability, not independent replication. At five neighbouring model settings, modularity and the connectivity–transition association held in every participant, and the correspondence with Movie10 exceeded its null in all but one fit; the association with mean first-passage time was less stable (**Table S10**). Assigning other recordings to *Friends*-defined states can miss patterns specific to them. Continuous-state and co-activation models (Lurie et al., 2020; Islam et al., 2024; Vidaurre et al., 2025) and independent fits to each experience would test how much of this organization depends on the discrete model.

## Conclusion

Across tens of hours of viewing, each person’s brain activity returned to a reproducible repertoire of states, and which state was expressed from moment to moment was related to the video, audio, and language content being viewed. That relation generalized to new films and, for language features, to reading and listening. The ordering of state use corresponded beyond what each recording’s power spectra and covariance reproduced in every participant for audiovisual film, but not consistently for reading or listening.

## Methods

### Participants and stimuli

Six healthy adult participants from the Courtois NeuroMod project (CNeuroMod, https://cneuromod.ca) viewed the television show *Friends* during longitudinal MRI scanning. The participants were aged 31-47 years old at recruitment in 2018. Three participants are women (sub-03, sub-04 and sub-06) and three are men (sub-01, sub-02 and sub-05). All participants reported right-hand dominance and good general health. Three were native French speakers (subjects 01, 02, and 04), one native English speaker (subject 06), and two bilingual native speakers (subjects 03 and 05). This distinction is relevant to the French/English *Le Petit Prince* comparison in the cross-stimulus analyses. All participants gave written informed consent under CNeuroMod’s ethics protocol. Data acquisition, demographics, and scanning logistics are documented in the official CNeuroMod documentation (https://docs.cneuromod.ca/en/latest/) and in the cited dataset descriptors.

Data collection took multiple scanning sessions across about three years, with each participant viewing the first six seasons of the American sitcom *Friends*, except sub-04, who watched the first four seasons. Most episodes were split into two ∼12 min scanner runs (labeled a and b) to accommodate participant comfort and minimize within-run head motion. Double episodes were split into up to four segments (a, b, c and d). The full *Friends* corpus thus contributed approximately 146 episodes (∼292 fMRI runs) per participant when all six seasons were completed. Participants viewed each episode passively (no task instructions, no overt responses required, no comprehension probes) through MR-compatible headphones (audio) and an MR-compatible projection display (video). We use *run* to refer to a single half-episode scanner acquisition and “episode” to refer to the full television episode (runs were concatenated only at the data-handling level, never during fitting).

We analysed every *Friends* run in the released CNeuroMod derivatives for seasons 1–6, except sub-04’s four season-5 runs (episodes 1 and 2), which were too few to form a season in the season-stratified split. The released data contain no run s05e20a for sub-03 and no runs s04e01a, s04e01b or s04e13b for sub-05. Two runs were present twice; we used one copy of each (sub-02 s05e09a: a 33-volume partial acquisition; sub-06 s02e04b: the first viewing). This left 292, 292, 291, 194, 289 and 292 runs for sub-01 to sub-06. We use the per-participant run count throughout the paper rather than aggregating across participants, and report all subsequent per-state and per-feature analyses with per-participant ranges (min–max across the six participants examined) rather than pooled summary statistics; the design treats N = 6 as six in-depth case studies rather than as a sample from a population.

In addition to *Friends*, all six participants also watched ∼10h of movies segmented into runs of ∼10min each while undergoing fMRI (the CNeuroMod *Movie10* dataset). Movies included three Hollywood feature films (*The Bourne Supremacy* (2004), *The Wolf of Wall Street* (2013), *Hidden Figures* (2016)) and one BBC nature documentary (*Life: Challenges of life, reptiles and amphibian mammals* (2009)). *Hidden Figures* and the *Life* documentary were both seen twice by each subject, totalling 61 runs per subject (*Bourne*: 10 runs; *Wolf*: 17 runs; *Life*: 2×5 runs; *Figures*: 2×12 runs). All participants except sub-04 also contributed to the *Harry Potter* dataset, for which they read chapter 9 of *Harry Potter and the Sorcerer’s Stone (1997)* inside the scanner. The text was presented word by word at a 2Hz pace (each word presented for 0.5s), and split across 7 fMRI runs of ∼9min to 14min administered in a single scanning session. The same five participants also listened to an audiobook version of *Le Petit Prince*, narrated in English and in French by a female-sounding voice, and split into 18 fMRI runs (9 runs per language) of about 9 to 13 min, acquired over three sessions; sub-06 contributed 16 runs (7 in French, 9 in English).

### fMRI preprocessing, parcellation, and dimensionality reduction

Throughout, our pipeline followed reproducible-neuroimaging practices consistent with the Standards recommendations from ReproNim (Kennedy et al., 2019): BIDS-organized inputs, preprocessing with recorded software versions, and standard cortical and subcortical atlases, with analysis code shared publicly. We used the fMRIPrep derivatives released by CNeuroMod (Esteban et al., 2019), produced with fMRIPrep 20.2.3, 20.2.5 or 20.2.6 depending on the *Friends* session, 20.2.3 for *Movie10*, 20.2.1 for *Harry Potter* and 20.2.6 for *Le Petit Prince*. The BOLD data were corrected for head motion and susceptibility distortion and resampled to the 91k CIFTI grayordinate space (fsLR 32k surfaces for cortex; MNI152NLin6Asym, 2 mm, for subcortical and cerebellar voxels). Slice-timing correction was applied to some of the *Friends* runs processed with fMRIPrep 20.2.3 (counts below and in **Table S7**). It was not applied to any *Movie10*, *Harry Potter* or *Le Petit Prince* run. We then applied a minimal-strategy confound regression (Ciric et al., 2017; Wang et al., 2024): 6 rigid-body motion parameters, the mean white-matter signal, the mean cerebrospinal fluid signal, and a discrete-cosine high-pass basis (cut-off 0.008 Hz), regressed out together with a linear trend (nilearn signal.clean); we applied no global-signal regression and no scrubbing. Each grayordinate’s time series was z-scored within run before parcellation, so each grayordinate contributes equally to its parent parcel’s mean.

The 1,650 analysed *Friends* runs were preprocessed with three fMRIPrep releases (20.2.3 for 1,056 runs, 20.2.5 for 529 and 20.2.6 for 65). Slice-timing correction was applied to 938 runs, all processed with 20.2.3; the other 118 runs processed with 20.2.3, all of them sub-01’s, were not corrected. Preprocessing changed with season in five of the six participants; all of sub-04’s runs used 20.2.3 with slice-timing correction. Reprocessing was not possible within the released derivatives.

In the 12 seasons that mixed preprocessing (14 contrasts in five participants), mean state occupancy did not differ detectably between runs with different preprocessing under an exact session-level permutation null (all p ≥ 0.12). Power was low, however, the minority preprocessing covered one to five sessions, and 7 of the 14 contrasts could not reach p < 0.05 by construction. Under a null that preserves acquisition order, two contrasts lay at the edge of the reference range (sub-02 and sub-05, season 5). In both, the difference traced to a single block of sessions rather than to the preprocessing groups as a whole. sub-06’s switch falls exactly at the boundary between seasons 4 and 5, so it could not be tested within a season. In the participants that could be tested, season-specific states are therefore unlikely to reflect the preprocessing change, though preprocessing and calendar time cannot be fully separated.

Preprocessed and z-scored grayordinate time series were aggregated into a 156-parcel composite atlas: 100 cortical parcels from the Schaefer Yeo 7-network atlas (Schaefer et al., 2018), plus a 56-parcel subcortical and cerebellar composite (CIT168 striatal/pallidal, Pauli et al. 2018; HCP thalamic, Glasser et al. 2016; HCP subcortical amygdala/hippocampus/brainstem; Cerebellum from the same HCP release). For network-level summaries we grouped the parcels into 13 networks: the seven Yeo networks of the cortical parcels and six subcortical groups (basal ganglia, 16 parcels; midbrain dopaminergic nuclei, 2; other midbrain and diencephalic nuclei, 8; thalamus, 14; hippocampus and amygdala, including the extended amygdala, 6; cerebellum, 10). The basal-ganglia group follows the circuit described by Alexander, DeLong and Strick (1986) and Haber and Knutson (2010). Each parcel’s time-series at each TR is the spatial mean of that parcel’s grayordinate-z-scored signals.

To reduce the parcel time series to a tractable subspace for HMM fitting, we applied principal-components analysis (PCA) per participant on a season-stratified training-split (70% train / 15% validation / 15% test), retaining the principal components needed to capture 95% of training-set variance. This yielded 67–77 PCs per participant. The variance threshold was part of the model-selection sweep (80%, 85%, 90%, 95% and 99%; see **Hidden Markov model**). The pipeline’s automated criterion, validation log-likelihood per PCA dimension, favored 99% in five of six participants, but it is biased toward more components because low-variance components add positive terms to the per-dimension likelihood, so we set 95% by hand. This keeps the component count (67–77) far below the number of time points. The reliability analyses below were run at 95% only; they show that this solution reproduces, not that 95% outperforms other thresholds. Per-component loading diagnostics, residual variance by parcel and network, and motion-artifact flags are reported in **Figure S2**. Our reliability analysis showed that the state set discovered from PCA with a 95% variance threshold reproduces under leave-one-season-out and split-half refits (see **Reliability**). Validation, test, and out-of-stimulus run data were projected through the training-fit PCA before any downstream modeling.

### Hidden Markov model

A hidden Markov model (HMM) treats the recording as a sequence of latent brain states that are not observed directly. At each TR t the brain occupies one of *K_max_* discrete states *z_t_*, and the observed activity *x_t_*, the D-dimensional PCA-projected parcel vector (D = 67-77 components; see **PCA**), is drawn from a distribution specific to that state. The states evolve as a first-order Markov chain, so the next state depends only on the current one. With π the initial-state distribution, A the *K_max_* × *K_max_*transition matrix (A_ij_ = P(z_t_ = j | z_t-1_ = i)), and Gaussian emissions x_t_ | z_t_ = k ∼ N(μ_k_, Σ_k_), the joint distribution of states and observations factorizes as

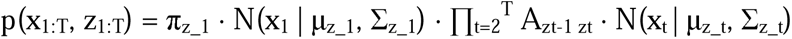

where N(· | μ_k_, Σ_k_) is the Gaussian density of state *k*. Each state is therefore a recurring brain activity pattern (its emission mean μ_k_) that the brain can re-enter at any time, while *A* and the dwell times it implies describe how those patterns follow one another. We chose an HMM because this structure matches our scientific target directly: it makes discrete state changes and their transition structure first-class objects of inference, supplying the dwell times, transition matrices, and Markovian timescales our questions require, and its states are latent and data-driven, inferred from the emissions rather than fixed in advance by stimulus events or experimenter labels. We fit a Gaussian-emission HMM by expectation-maximization over this finite state space of fixed maximum capacity *K_max_*. Onto its transitions we place two priors taken from the sticky hierarchical Dirichlet process HMM (sHDP-HMM, Fox et al. 2011): a sticky bias on self-transitions that discourages single-TR state switching and favors multi-TR dwell, keeping state durations compatible with the slow timescale of the hemodynamic response; and a hierarchical Dirichlet concentration prior on the transition rows that shrinks occupancy toward fewer states. We borrow this prior structure but not the sHDP-HMM’s nonparametric inference: we never infer the state count over an unbounded space. We fix *K_max_* and let the number of occupied states emerge from the data below that ceiling. With the concentration prior at its production setting (γ = 1) the prior alone is parsimonious: a fit-free prior-predictive simulation that draws transition matrices from exactly this prior occupies a median of 2 states above a 1% usage threshold (95% interval 1–6), far fewer than the data occupy.

We fit one combined model per participant across that participant’s PCA-projected runs, so a given state index refers to the same emission distribution wherever it appears and no post-hoc matching of states across episodes is required. Fitting is by expectation-maximization under a finite (weak-limit) truncation of the prior: the M-step updates each state’s Gaussian emissions (its mean and a diagonal covariance in PC space) by maximum likelihood, regularized only by a variance floor of 10^-3^, and the transition rows are updated by their posterior-mean (Bayesian point) estimate under the sticky, hierarchical-Dirichlet prior. The shared state weights β that set the prior mean of every row are re-estimated at each iteration from the summed transition counts. The estimator is therefore maximum-likelihood on the emissions and posterior-mean on the transition rows. The concentration hyperparameters (γ, α) and the sticky bias (κ, ρ) are held fixed, all *K_max_* states are retained (none are pruned), and we decode state sequences with the Viterbi algorithm. As in other fMRI HMM studies (Vidaurre et al., 2017, 2018; Baldassano et al., 2017), the number of states is fixed before fitting rather than inferred. The truncation capacity *K_max_*is a fixed ceiling, chosen from the sweep described below and high enough not to be reached: under the final configuration 37–42 of the 50 states exceed 1% usage and 42–47 are active. Each state’s emission covariance is diagonal in PC space, so the model does not fit covariance between principal components within a state. Because principal components are weighted mixtures of parcels, a covariance that is diagonal in PC space is not diagonal in parcel space: projected back to parcels, each state carries a full covariance, so the diagonal choice does not assume that parcels are uncoupled within a state. We measure coupling between parcels within a state directly in a separate analysis (see **State transition structure**) rather than reading it from the PC-space covariance.

We chose the final configuration (*K_max_* = 50; γ = 1; κ = 10; α = 1; ρ = 1) from a sweep fitted on the training split and scored on the validation split, with five initializations per configuration. The sweep crossed *K_max_* ∈ {20, 40, 50, 60, 80, 100} with γ ∈ {1, 5, 10} at variance thresholds of 90%, 95% and 99% (κ = 10, α = 1, ρ = 1 and diagonal covariance throughout), and added 80% and 85% at *K_max_*= 60, γ = 5. For each participant we compared validation log-likelihood with the number of occupied states, a state counting as occupied when its usage fraction at the final fitting iteration exceeds 1% (**Figure S3**). We set one configuration for all participants by hand rather than taking the pipeline’s automated choice (99% variance and *K_max_* = 100 in most participants). We did not select by likelihood alone, because likelihood-based criteria (validation log-likelihood and BIC) favor the highest *K_max_*: added capacity is absorbed by states the data barely visit, which still raise the likelihood, while the number of meaningfully occupied states stays bounded. The occupied-state counts fell into two groups. With γ = 1, every capacity from *K_max_* = 40 to 100 gave 35–44 occupied states in every participant; with γ = 5 or 10, capacities of 60 and 80 gave 50–62 (45–56 at 100), adding states for little gain in validation log-likelihood and with a larger train-to-validation gap. Within the lower group, *K_max_* = 40 filled 88–98% of its capacity (35–39 of 40 states occupied), so a count close to the ceiling cannot be told apart from one limited by it. At *K_max_* = 50 with γ = 1, 36–41 of the 50 states were occupied, leaving room for states the fit did not use. We therefore took *K_max_* = 50, the smallest capacity whose occupied count stayed clear of the ceiling, and the lowest concentration, γ = 1: a prior that favours fewer states and still finds this many is stronger evidence that the data support them. The chosen configuration lies on the Pareto front of validation log-likelihood against occupied states in four participants; in sub-01 and sub-02 a γ = 1 model with higher capacity has slightly higher validation log-likelihood with no more occupied states. Its train-to-validation gap is smaller than that of every γ = 5 or 10 model with higher capacity. Under this configuration the usage-based count is 37–42 states per participant. The repertoire carried into the downstream analyses is defined by recurrence rather than usage (a state is active if its per-run fractional occupancy exceeds the activity threshold in at least one run; see **State classification**) and comprises 42–47 states per participant.

The selected configuration, the same for all six participants, was refit for each participant with 10 random initializations on the combined train-plus-validation split; we kept the initialization with the highest combined train-plus-validation log-likelihood as the primary model. We then decoded every run (train, validation and test) as a separate sequence with the Viterbi algorithm, which can assign any of the 50 states; the active repertoire is defined from these sequences (see **State classification**). We assessed within-*Friends* reproducibility of this repertoire with leave-one-season-out and split-half refits (see **Reliability**).

### Reliability

Each participant’s primary HMM was fit on that participant’s *Friends* training and validation runs (85% of runs; the **PCA** on the 70% training split) and decoded on all runs, leaving open whether the discovered repertoire reproduces under independent fits or remains specific to the exact runs included in training. To assess within-*Friends* reproducibility, we ran two refit procedures that control different confounds: leave-one-season-out (LOSO) refits test whether the repertoire depends on the specific seasons used for fitting, and split-half refits test whether the repertoire reproduces across interleaved odd/even episode halves and so reduces the session-order confound that LOSO does not isolate (session-order spans the full episode index whereas individual seasons are partly correlated with session-position). The two procedures together characterize different facets of within-*Friends* reproducibility, stability under removal of a season (LOSO) and order-independence (split-half).

For LOSO, we partitioned each participant’s seasons into one fold per available season, 34 folds in total (six folds for sub-01, sub-02, sub-03, sub-05, sub-06; four folds for sub-04, who contributed Seasons 1–4 only; see **Participants**). Each fold held out one season’s runs from both PCA fitting and HMM fitting, refit under the primary configuration on the held-in seasons, and produced a Viterbi-decoded state sequence for the held-out runs. For split-half, each participant’s episodes were partitioned into two halves by an interleaved odd/even rule on episode order within each season; each half refit its own PCA and HMM independently under the same configuration, with the two halves’ models compared post-hoc by matching their state indices.

Each LOSO fold and each split-half fits its own PCA on the appropriate training subset. Reusing the primary model’s PCA, whose training split contains runs from every season and from both halves, would leak held-out data into the per-fit basis, because the principal components themselves would be informed by the held-out runs’ covariance. Refitting per fold isolates the train/held-out partition at the level of the basis as well as the HMM.

Each refit produces an independently-indexed *K_active_*state set whose state labels are not directly comparable to the primary model’s. Because each fit learns its Gaussian emission means in its own PCA subspace, we back-projected every state’s mean to the common 156-parcel space before matching. For state k with PC-space emission mean μ_k_ ∈ □^P^ over the P retained components, the mean parcel-space activation vector is m_k_ = W□ μ_k_ + c, where W ∈ □^(P×N)^ stacks the P retained principal-component loadings (the first P rows of the PCA components matrix), c ∈ □^N^ is the training-set parcel mean, and *N* = 156 (P is the fit-specific component count: 67–77 for the primary models and 66–78 across the LOSO and split-half refits; see **PCA**). We then matched states post hoc by the Hungarian algorithm (Kuhn 1955; Munkres 1957) on the matrix of Pearson correlations between the two fits’ active-state vectors (*K_A_* × *K_B_*; active meaning fractional occupancy above 0.02 in at least one decoded run), returning the one-to-one pairing of min(*K_A_*, *K_B_*) states that maximizes the summed paired correlation. Matched pairs with r > 0.3 were treated as well-matched. This cutoff is a permissive screen rather than the reproducibility evidence itself: because the Hungarian algorithm returns a one-to-one pairing for every state, including states with no genuine counterpart in the other fit, the threshold serves only to separate substantive matches from forced assignments. We set it low, at a modest correlation by conventional benchmarks, so that it flags spurious pairs without imposing a stringent bar that would itself shape the well-matched count. The actual evidence is the full matched-pair correlation distribution (**Figure S4**), which lies well above this floor.

Hungarian matching on parcel-space does not test whether matched states share within-state functional-connectivity (FC) structure: two states with similar mean activation patterns but different FC would be falsely matched. To address this caveat, we additionally computed empirical within-state FC for every LOSO fold and split-half: for each state we pooled the TRs assigned to it across the runs that fit decoded (for a LOSO fold, the held-out season; for a split half, all of its runs) and computed a Ledoit–Wolf-shrinkage Pearson correlation matrix from the parcel time series; each LOSO fold’s matrices were compared with the primary model’s, computed over all runs. We used shrinkage because a state’s FC is estimated only from its assigned TRs (a median of about 500 per state in a LOSO held-out season, and fewer than the 156 parcels for 5–12% of states); the sample covariance is then poorly conditioned and noisy, whereas Ledoit–Wolf shrinkage toward a scaled-identity target gives a well-conditioned, lower-variance estimate with a closed-form optimal intensity (Ledoit & Wolf, 2004). States with fewer than 30 assigned TRs were flagged; the pairs that involve them (0–9 per participant and refit type) are kept in **Figure S4**, and dropping them does not change the comparison. For each matched pair we then computed the RV coefficient between the two FC matrices (Robert & Escoufier, 1976), a correlation-like similarity index for two matrices that ranges from 0 (no shared structure) to 1 (identical up to scale), defined as RV(A, B) = tr(AB) / √(tr(A²)·tr(B²)). Because every within-state FC matrix is dominated by a component shared across all states, the RV coefficient computed on the raw matrices is close to its ceiling whether or not two states correspond: matched pairs and deliberately mismatched pairs differ by 0.028 or less (0.003–0.028 across participants and refit types). Raw RV therefore cannot flag a mean-only match, and we do not use it for that purpose. We instead subtract each fit’s across-state mean FC, compute the same trace ratio on the residual matrices without clipping it to [0, 1] (residuals can be negatively similar), and compare it with a mismatched-pair null in which each state is paired with a different state of the other fit (one derangement of the matched indices). On that measure the matched mean exceeds the null mean in every participant and refit type, by 0.09–0.58; the smallest gap is in sub-02’s split halves (0.23 against 0.14), and every other gap exceeds 0.27. Matched states therefore share connectivity structure beyond the component common to all states (**Figure S4**). The comparison is descriptive; no test statistic was computed..

We report four reproducibility metrics: the matched-pair correlation distribution (each LOSO fold against the primary model, and split half A against half B); per-fit structural invariants (the number of states the fitted model uses, *K_used_*, defined by posterior usage above 0.01; transition entropy and self-transition probability over those states; and the median and interquartile range of Viterbi dwell time), compared as scalars across folds and between halves; for split-half, the Spearman correlation between the recurrence scores of matched states with r ≥ 0.3 (testing whether the rank ordering of states by recurrence persists across independent fits); and Kolmogorov–Smirnov tests comparing the distribution of per-state recurrence scores between every pair of LOSO folds (Benjamini–Hochberg-corrected within participant) and between the two split halves. The matched-pair correlation distributions and the LOSO structural invariants are shown in **Figure S4**; the split-half recurrence ordering is shown in **Figure 1D**. To contextualize the scale of fit-to-fit variability we compared the LOSO fold-to-fold spread of *K_used_*, transition entropy and self-transition probability with their range across the 10 random-seed initializations of the primary model, a relative-scale anchor only, not a confidence interval; median dwell time has no seed comparison. Within-*Friends* reproducibility evidence is reported per participant.

To check whether the states depend on the model’s temporal assumptions, we compared them with spatial independent component analysis (ICA), which assumes spatially independent sources and has no temporal model. At each dimensionality K (15, 25, 35 and 41–47 components, and each participant’s *K_used_*), ICA was fitted in the subspace of the top K principal components of the same PCA (FastICA, 25 restarts combined into consensus maps). ICA maps were matched one-to-one to the content-eligible states’ mean maps by absolute spatial correlation, and the median matched correlation was compared with that of 1,000 random rotations of the same whitened top-K subspace, with the matching rerun for each rotation (one-sided empirical p per participant and K). ICA matched the state maps better than these rotations, but only slightly (median |r| higher by 0.02–0.11; p < 0.05 in 56 of the 60 participant-by-dimensionality cells of **Figure S10A**; a 57th, sub-06 at K = 15, lies at the threshold, p = 0.051), and did not reproduce the states one by one; most of the agreement came from the shared subspace. We also applied each participant’s *Friends* ICA components at *K_used_*(37–42) to the other stimuli without refitting and correlated their *Friends* recurrence with their occupancy, as in the cross-stimulus analysis but against 1,000 phase-randomized surrogates (**Figure S12**). The correlation exceeded its phase-randomized null for *Movie10* in all six participants, for *Le Petit Prince* in all five and for *Harry Potter* in two. Because winner-take-all component labels impose no minimum visit length, most components exceed the 2% occupancy threshold in most runs (median recurrence 0.64–0.81, against 0.45–0.51 for the states), so ICA recurrence is not comparable with state recurrence and these correlations do not test the main analysis.

### State classification: recurrence, season specificity, and taxonomy

From each participant’s decoded state sequence, we computed fractional occupancy (FO) per state per run (the proportion of a run’s TRs spent in each state) and derived a per-state recurrence score: the fraction of that participant’s runs in which the state’s FO exceeded a 0.02 activity threshold (about 9 TRs, or 14 s, at the mean run length of 472 TRs). In sub-02’s 33-volume partial run of s05e09a (see **Participants and stimuli**) the threshold is below one TR, and one state is active only in that run, so sub-02’s active repertoire is 46 states rather than 45. Recurrence is a continuous score in [0, 1]. States with recurrence > 0 (occupied above threshold in at least one run) form each participant’s active repertoire, 42–47 states, the set carried into the downstream analyses. Downstream analyses work from this continuous score rather than from a binary recurrent/non-recurrent label: the screening taxonomy thresholds it, and the transition and cross-stimulus analyses use it as a per-state value.

To identify states whose occupancy concentrates in particular seasons rather than spreading evenly across the participant’s viewing history, we computed a season-specificity index (the range, maximum minus minimum, of the state’s per-season recurrence scores; 0 when recurrence is the same in every season, 1 when the state is active in every run of one season and in no run of another). We tested the season-specificity index against a null generated by shuffling season labels 5,000 times within participants, with the per-state permutation p-value computed as (count + 1) / (n_permutations + 1). We applied Benjamini–Hochberg correction (Benjamini & Hochberg, 1995; *q* < 0.05) within each participant separately to control the per-state false-discovery rate, and treated the FDR-corrected significant set per participant as that participant’s season-specific states. The analysis treats *N* = 6 as six in-depth case studies and does not pool *p*-values across participants.

To partition the recurring repertoire into categories that expose the identifiable sources of recurrence, we synthesized four families of per-state diagnostic flags into a single mutually-exclusive taxonomy. States with a recurrence score of 0 were never assigned by the decoder and were excluded before categorization. The remaining states form the active repertoire. Each active state was assigned to one of five mutually exclusive categories by descending priority: (1) low-confidence (sub-HRF; median dwell < 3 TRs), (2) run-onset-anchored (visits concentrated in the first fifth of runs), (3) drift-anchored (fractional occupancy varying with season and trending with global session position), (4) content-eligible (no exclusion tag and recurrence ≥ 0.10), and (5) rare (no exclusion tag but recurrence < 0.10). A state flagged by more than one criterion is reported under the highest-priority match. For example, a sub-HRF state active in fewer than 10% of runs is reported as low-confidence rather than rare, because signal quality takes precedence; sub-HRF states were not tested for run-onset anchoring. These screens use only temporal information (usage, dwell time, run-onset timing and session-position drift) and do not inspect a state’s spatial map. Content-eligible is therefore a residual label: it marks states with no identified temporal confound and does not rule out spatially artifactual sources. Downstream analyses use this taxonomy in two ways. Transformer-depth analyses use only content-eligible states. Cross-stimulus recurrence-transfer analyses use the full active repertoire as the primary state set and repeat the correlations on the content-eligible subset as a robustness check.

### Network participation of recurring states

To describe what recurred spatially, we summarized each content-eligible state’s parcel-space mean map against the 13 network labels of the parcellation (the seven Yeo cortical networks and six subcortical groups: basal ganglia, dopaminergic midbrain, midbrain–diencephalic, thalamus, hippocampus/amygdala and cerebellum; see **fMRI preprocessing, parcellation, and dimensionality reduction**). For each state we took the absolute value of each parcel in its mean map and averaged these within each network label (averaging across a network’s parcels, so that larger networks are not mechanically favored), giving a per-network participation magnitude; because the metric uses absolute loadings, it reflects participation magnitude and does not separate activation from deactivation. Normalizing the per-network values to sum to one gave each state’s network composition, from which we computed the largest-network share, the summed top-three share, the number of networks contributing at least 10% of the composition, and the network-spread index (the normalized Shannon entropy of the composition: its Shannon entropy divided by the log 13 (the number of network labels); 0 when all magnitude falls in one network and near 1 when it spreads evenly across labels). We summarized these metrics across each participant’s content-eligible states and across the cohort and reported the most common top-three network sets. **Figure 2B** and **Figure S11** apply the same metrics to every active state (269 across participants), grouped by category; **Figure S11** shows all 13 networks.

### Matching states across participants

For each of the 15 participant pairs from 6 participants, we correlated every content-eligible state map of one participant with every map of the other. Maps were defined as in Joint embedding (state mean relative to the participant’s mean, z-scored across regions) and compared with Pearson correlation. We then paired states one-to-one to maximise the summed correlation (Hungarian algorithm), so that every state of the participant with fewer states received a partner. The pair statistic was the median correlation over matched states.

To test whether matches exceeded what two repertoires would give by chance, we compared this statistic with a state-mixing null. In each of 1,000 draws, one participant’s centred state maps were mixed by a random orthogonal matrix (alternating between the two participants), the maps were re-z-scored, and the matching was recomputed. The mixing keeps each participant’s mean map and the full spatial covariance of their maps, including any dominant spatial patterns shared across participants, but destroys which state is which. One-sided *p* values were computed as (1 + number of null values ≥ observed) / (1 + number of draws), and corrected across the 15 pairs with the Benjamini–Hochberg procedure.

We report two specificity tests; each version of the maps had its own state-mixing null. First, to ask whether matches reflected more than shared large-scale network composition, we replaced each region’s value with the mean of its network or subcortical structure (13 groups, so that each group’s mean entered the correlation once per region), and separately with the complement (the map minus those means, i.e. the pattern within networks). Second, to ask whether matches reflected only components common to everyone, we projected out of both participants’ maps the three largest components of a PCA over the other four participants’ state maps.

To ask whether matched states also occurred at the same moments of the stimulus, we correlated the binary occupancy time courses of each matched pair over the runs both participants watched (the same run identifier denotes the same segment of the same episode). Runs were aligned from their start and cut to the shorter run. The null circularly shifted one participant’s occupancy within each run (1,000 shifts). At the pair level we compared the mean locking correlation over matches with this null. Individual matches were flagged as time-locked when their correlation exceeded a family-wise threshold, the 95th percentile of the null distribution of the maximum over the pair’s matches. For comparison with a ceiling, we applied the same matching to two models fitted separately to halves of each participant’s data. All inference is per participant pair. Individual map matches are descriptive, because a per-match threshold on the state-mixing null was liberal in calibration.

For each match we correlated the two states’ binary occupancy time courses over the runs both participants watched (runs aligned from their start and cut to the shorter run). The second participant’s time course was shifted by −20 to +20 TRs (about ±30 s) within each run, without wrapping. All 348 matches were included, not only the time-locked ones, so a peak at zero lag is not produced by the selection. The zero-lag value equals the occupancy correlation used in the time-locking test (**Figure S14E**).

### Joint embedding of participants’ states

We embedded participants’ activity maps in two dimensions with UMAP (umap-learn 0.5.12; McInnes et al., 2018). Each state was represented by the same map used for cross-participant matching: its mean activity across the 156 regions relative to the participant’s mean (the state mean minus the participant’s PCA mean), z-scored across regions. The maps of all six participants were embedded together using correlation distance, 15 neighbours, a minimum distance of 0.3 and a fixed random seed. We computed two embeddings: one of the content-eligible states (n = 158; **Figure 2E**, **Figure S14A, C**) and one of all active states (n = 269; **Figure S14B**). In **Figure 2E**, lines join the one-to-one matches between each pair of participants (Hungarian matching on map correlation; see **Matching states across participants**). Line width and opacity scale with the matched correlation, and dark lines mark matches flagged as time-locked. **Figure S14A** shows the same embedding with the matches recomputed after the three largest components of a PCA over the other four participants’ state maps were projected out of both participants’ maps. The embedding is a display only: distances and clusters in it were not tested, and all inference on cross-participant correspondence is per participant pair. To check that the layout does not depend on the UMAP settings, we re-embedded the content-eligible states with 5, 15 and 30 neighbours at three seeds each, and with PCA and classical multidimensional scaling of the correlation distances (**Figure S14D**).

### State transition structure

We described each participant’s transition structure from two sources. (1) The model’s learned transition matrix, which expresses the long-run dynamics with the sticky self-transition prior made explicit. It was used for the display in **Figure 3A** and for the module partition shown there, which also defined the modules in the analyses of module dwell, module timing and co-membership; for the predicted-matrix test; for the stay probabilities and destination popularity of the chain null; and for the mean first-passage time. (2) Empirical transition counts tallied from the Viterbi-decoded sequences within runs. They were used for the modularity tests, the functional-connectivity (FC)–transition Mantel test, network homophily and recurrence assortativity. Module dwell and timing were measured on the decoded sequences themselves. All permutation p-values were computed as (1 + number of null values ≥ observed) / (1 + number of draws) (Phipson & Smyth, 2010). They were one-sided unless stated otherwise; two-sided p-values compared the absolute values of the null and observed statistics.

State-pair functional similarity was the RV coefficient (Robert & Escoufier, 1976) between the two states’ within-state FC matrices. Within-state FC is the pattern of co-activation among parcels while a state is active; each matrix was a Ledoit–Wolf shrinkage correlation over the TRs assigned to that state. FC was not estimated for states with fewer than 30 assigned TRs (one state each in sub-01 and sub-02 and two in sub-05, each active in a single run). These states were left out of the predicted-matrix test and the co-membership analyses, and their pairs were treated as missing in the Mantel and MFPT tests. Each state was assigned a dominant network among the Yeo 7 networks and 6 subcortical groups (basal ganglia, dopaminergic midbrain, midbrain–diencephalic, thalamus, hippocampus/amygdala, and cerebellum) by maximum mean absolute activation.

We tested whether each participant’s transitions grouped states into modules using directed weighted modularity Q (Leicht & Newman, 2008). Q compares the transition probability within groups of states with what each state’s overall rates of leaving and entering predict. We maximized Q over partitions with the Louvain algorithm (Blondel et al., 2008) at resolution 1, ran it from 50 fixed seeds and kept the best partition. Every observed and null graph was treated the same way. The nodes of the tested graph were the active states. Its edges were the transition probabilities counted from the Viterbi-decoded sequences within runs, normalized over all next states including the current one, as in the model’s matrix, with self-transitions then removed; edges with probability below 0.01 were dropped (296–364 directed edges per participant).

We compared each graph with two nulls that keep a simpler structure. The shuffled-order null shuffled the order of decoded segments within each run. This kept which states occurred in each run, the length of every segment and the run boundaries (1,000 draws). The chain null simulated a Markov chain without modules. It kept each state’s fitted probability of staying and its popularity as a destination, along with the observed run lengths and starting states (500 draws). P-values were one-sided, (1 + number of null values ≥ observed) / (1 + number of draws). A participant’s transitions were called modular when three conditions held: both nulls gave *p* < 0.05; Q exceeded the mean of the shuffled-order null by at least 0.05; and the median adjusted Rand index between the observed partition and those of 200 run-level bootstrap resamples was at least 0.5.

We checked the nulls on simulated chains. The shuffled-order null rejected none of 200 chains without modules, and all 200 chains with planted modules at each of the two module strengths tested. A null that rewired edges while preserving degree and reciprocity, with weights matched to node strength, rejected 38.5–42.5% of graphs without modules, so we did not use it for inference. For each leave-one-season-out fold, the held-out season was decoded by that fold’s model, and its counted transitions were tested against the shuffled-order null. The partition shown in **Figure 3A** is that of the learned transition matrix, thresholded at the same 0.01 (335–398 directed edges per participant), which removes weak and likely spurious edges. All participants, graph variants and nulls are reported in **Table S6**.

Module dwell was the mean number of consecutive TRs spent within one module, compared with the same quantity in the shuffled-order sequences (1,000 draws). For cross-participant timing, each module’s activity map was the mean of its states’ maps, weighted by how many TRs each state occupied. Each participant’s modules were then matched one-to-one to another participant’s by Hungarian matching on the correlation of these maps. For each matched pair of modules, we marked at every TR whether each participant was in that module, and correlated these indicators over the runs both participants watched, truncating each run to the shorter of the two. The statistic was the mean correlation over matched pairs. Its null came from 1,000 draws in which each of one participant’s runs was circularly shifted by a random offset.

We used seven pairwise measures that do not depend on the order in which states were visited: (1) functional similarity; (2) the correlation of activation maps; (3) the Bhattacharyya distance between the states’ emission distributions; (4) whether the states shared a dominant network; (5) the absolute difference in recurrence; and (6) co-occurrence across runs (the Spearman correlation of the two states’ occupancy over runs) and (7) co-occurrence across seasons (the Pearson correlation of the two states’ mean occupancy in each season). To ask whether the modules exceeded what these relations predict, we regressed the log off-diagonal transition probabilities of the model’s matrix on the seven measures. The regression also included the log out-strength of the source state and the log in-strength of the target state. Each null matrix was the fitted prediction plus the residuals permuted across state pairs, thresholded at 0.01; the best Q of 1,000 such matrices formed the null. To describe which relations the modules followed, we compared the mean of each measure between pairs of states in the same module and pairs in different modules. Its null permuted module labels across states, which keeps module sizes (5,000 permutations, two-sided). We then entered all seven measures jointly in a multiple-regression quadratic assignment model of module co-membership (Dekker et al., 2007), with 5,000 permutations of state labels (two-sided).

The FC–transition Mantel test (a permutation test for association between two matrices) correlated, with Spearman’s ρ, the upper triangles of the state-by-state FC-similarity matrix and the direction-symmetrized empirical transition matrix. The null was 5,000 state-label permutations (two-sided), with the missing-cell mask re-intersected on each permuted matrix. FC profiles and transition counts both derive from the same Viterbi assignments. This null therefore tests whether their pairing within a fit is non-random, not the pairing of two independent measurements.

Mean first-passage time (MFPT), the expected number of TRs for a path from state i to first reach state j, was computed from the learned transition matrix. It was computed over the largest strongly connected component (the set of states all mutually reachable through transitions), which contained every active state in all six participants. We tested whether symmetrized MFPT distance co-varied with FC dissimilarity (1 − RV) under the same two-sided state-label permutation framework as the Mantel test.

To test whether transitions preferentially link states that share a dominant network, we contrasted the average transition probability of within-network state pairs with that of between-network pairs, each taken over the ordered state pairs with at least one observed transition. The test statistic was the within-minus-between difference, which avoids the division by zero a ratio incurs when the between-network probability approaches zero. It was evaluated against a null of 5,000 permutations of network labels across states. We report the effect size as the within/between ratio.

Recurrence assortativity, the tendency for states with similar recurrence scores to transition into one another, was Newman (2003)’s numeric assortativity coefficient, with recurrence score as the node attribute and each directed edge counted once, without its weight. It was computed on a denser graph than the one used for modules (transition probability ≥ 0.005; 626–724 edges) for greater statistical power. The null re-paired recurrence labels across nodes (5,000 permutations, two-sided). Recurrence was nearly a rank transform of each state’s long-run occupancy under the fitted transition matrix (Spearman’s ρ = 0.97–0.98 with the stationary distribution). States with high stationary probability are the target of more of these edges, and this null breaks that link. Assortativity measured this way therefore describes the transition structure, not an association between two independently measured properties.

### Cross-stimulus decoding and recurrence correspondence

To test whether the brain-state repertoire discovered from *Friends* carries to other naturalistic stimuli, we applied each participant’s *Friends*-fit pipeline (PCA + HMM) unchanged to three additional CNeuroMod stimulus datasets, chosen to span a modality gradient: audiovisual films (*Movie10*), word-by-word visual reading (*Harry Potter*), and audio-only narrative listening (*Le Petit Prince* in French and English). No part of the pipeline was fine-tuned, updated, or re-fit on the out-of-stimulus data; we ask only whether the *Friends*-defined states and their recurrence ordering correspond to what these stimuli evoke.

The three additional stimulus families differ in modality, run count, and per-participant coverage. *Movie10* comprises three feature films and one nature documentary (CNeuroMod release, Boyle et al., 2023) presented audiovisually, with all six participants completing 61 runs each. *Harry Potter* (HP) presents a chapter of *Harry Potter and the Sorcerer’s Stone* word-by-word via rapid serial visual presentation (Wehbe et al., 2014); five participants completed seven HP runs each (sub-04 does not appear in the HP dataset, a CNeuroMod data-provenance limitation, not a QC exclusion). *Le Petit Prince* (PP) presents continuous audiobook narration in French and English (stimulus and word annotations from Li et al., 2022); the same five participants listened to both languages, all nine sections of each except for sub-06, for whom seven French sections are available (16 runs). All four stimulus families, and the resting-state runs, were acquired on the same scanner with the same multiband EPI sequence (Siemens Prisma Fit 3 T; multiband factor 4; TR = 1.49 s; TE = 37 ms).

For each participant and each out-of-stimulus run, parcel time-series were projected through the participant’s *Friends*-trained PCA basis (vt = 0.95, 67–77 PCs per participant) without any re-fitting. The forward design preserves the one-to-one correspondence between *Friends* state indices and out-of-stimulus state indices by construction; refitting PCA on out-of-stimulus data would change the basis and forbid direct comparison of state assignments across stimuli. As a diagnostic of subspace alignment, reconstruction *R²* (the fraction of out-of-stimulus parcel variance recovered by the *Friends* PCA basis, centered by the *Friends* training-set mean) ranged 0.93–0.95 across the 16 (participant × out-of-stimulus dataset) cells. The *Friends*-trained subspace therefore captures most of the variance in out-of-stimulus parcel data. By network, *R²* was 0.92–0.98 in the six cortical networks other than the limbic network (0.62–0.86), 0.84 or higher in the basal ganglia, midbrain and hippocampus/amygdala groups, 0.70–0.91 in the cerebellum, and lowest in the thalamus (0.23–0.55), so the thalamic part of each state is the part the *Friends* subspace represents least well in the other stimuli.

The participant’s *Friends*-fit HMM (see **Hidden Markov model**) was applied to the projected PC time-series without any re-fitting or fine-tuning. Per-TR Viterbi decoding yielded the most probable state sequence given the fitted emission and transition parameters. Viterbi assigns every TR to one of the model’s states by construction, almost always a state that was active during *Friends* (at most 1.5% of any run’s TRs fell in states unused during *Friends*, and those states are left out of the correlations); the cross-stimulus test asks whether the *Friends* repertoire accounts for out-of-stimulus activity. We computed per-run fractional occupancy (FO) per state from the Viterbi sequence, as in the within-*Friends* analyses (see **State classification**); these FO correlations are the cross-stimulus tests.

For each participant, we computed Spearman’s ρ between the per-state *Friends* recurrence score (see **State classification**) and the per-state mean fractional occupancy across out-of-stimulus runs. The correlation was computed and tested separately per participant; we do not pool *p* values or test ρ at the group level, and the across-participant means of ρ given in **Results** are descriptive. We computed per-stimulus-subset variants in parallel: for *Movie10*, per-film breakdowns (*Bourne* / *Wolf* / *Figures* / *Life*); for *Le Petit Prince*, per-language breakdowns (French / English) within participants. Both the full-set and eligible-subset versions were computed; the full set is the primary cross-stimulus test and the eligible subset is reported as a robustness reference.

We ran each cross-stimulus correlation on two state sets. The primary analysis uses the full active repertoire (every state with recurrence > 0), because recurrence is a property of the whole repertoire rather than of any one subset. As a robustness check we re-ran it on the content-eligible states alone (the states left after the screening taxonomy removes the sub-HRF, run-onset-anchored, drift-anchored, and rare states; see **State classification**), testing whether the correspondence survives once those screened states are dropped. The content-eligible correlations were not tested against the phase-randomized null described below and are reported descriptively (**Table S3**).

To assess how much of the cross-stimulus correlation could arise from shared low-level temporal structure, we compared each cross-stimulus correlation (*Movie10*, *Harry Potter*, and *Le Petit Prince* with its French and English runs pooled) with a phase-randomized surrogate null (Prichard & Theiler, 1994). For each surrogate, we Fourier-transformed each of that stimulus’s runs’ projected PC-score matrices along time and applied a random phase shift at each frequency, using the same shift across all principal components. The zero-frequency (DC) term, which determines the run mean, and, for even-length runs, the Nyquist term, the highest frequency supported by the sampling rate, were left unchanged because both must remain real-valued to produce a real-valued time series after the inverse transform. This preserves each component’s power spectrum, variance, autocorrelation, and cross-component covariance, as well as the number and length of runs, while disrupting stimulus-locked phase alignment and higher-order temporal structure. Only the held-out stimulus’s PC scores were randomized; the *Friends* recurrence scores were kept fixed. For each participant and stimulus, we generated 10,000 surrogates, repeated Viterbi decoding, recomputed state occupancy and Spearman’s ρ, and compared the observed ρ with the surrogate distribution. Surrogates were deterministically seeded by draw index. Surrogates were decoded with a standalone diagonal-Gaussian Viterbi decoder rebuilt from the saved model parameters. Before any surrogate was drawn, this decoder had to reproduce each participant’s published ρ on the original runs (to within 10^-^¹²; it did so exactly for every participant and stimulus), so the null tests the same statistic as the original analysis. Significance was assessed using a one-sided empirical *p* value with the Phipson–Smyth correction (minimum *p* = 0.0001), with inference performed separately for each participant. Because phase randomization removes phase-dependent structure broadly rather than specifically stimulus locking, exceeding this null shows that the preserved spectra and covariance cannot fully explain the correlation, but does not by itself establish that the remaining correlation reflects stimulus-locked dynamics. We report z, the distance of the observed correlation from the null mean in null standard deviations, as a descriptive position; the empirical *p* is the inferential statistic. Each null pools a stimulus’s runs (61 for *Movie10*, 7 for *Harry Potter*, 16–18 for *Le Petit Prince*), so for *Movie10* the four films were not tested separately.

As a further test of whether recurrence relates to ongoing experience, we applied the same frozen pipeline to resting-state runs, which carry no external content stream. All six participants contributed resting-state data (four to six runs each), as for *Movie10*; *Harry Potter* and *Le Petit Prince* lack sub-04. Parcel time-series were projected through the participant’s *Friends*-trained PCA basis and decoded by the participant’s *Friends*-fit HMM, with no re-fitting or fine-tuning at either step, exactly as for the three narrative stimulus families. We computed per-run fractional occupancy per state and correlated it (Spearman’s ρ) with the per-state *Friends* recurrence score, again per participant without pooling. Two diagnostics accompany the correlation: reconstruction *R²* for the *Friends* PCA basis applied to rest, and the log-likelihood gap between held-out *Friends* runs and rest runs, which indexes how well the fitted temporal model accounts for resting dynamics. No surrogate null was run for the resting-state condition, so all resting-state values are descriptive and we do not test them against a null. Because raw ρ carries a stimulus-independent covariance and stationarity floor, we do not interpret the ordering of ρ among the weakly corresponding conditions as a specificity gradient. Resting-state results are shown in **Figure S8** and are not part of the main cross-stimulus figure (**Figure 4**).

### Transformer features and brain-state correspondence

We decoded the fitted state, and did not predict parcel activity, because the question was whether the states, estimated without the stimulus, were related to stimulus content. An encoding model tests whether content predicts regional activity; it does not test whether the recurring states identified by the HMM are related to content. State decoding should succeed to the degree that states summarize content-related activity. Decoding states also allowed a test on independent stimuli: whether a mapping from content to states learned on *Friends* held for unseen films, reading and listening.

To probe correspondence between brain states and stimulus content across the four stimulus families (*Friends*, *Movie10*, *Harry Potter*, *Le Petit Prince* in French and English), we extracted layer-wise activations from three frozen pretrained transformers, one per modality. For audio and text we used two of the three backbones validated in TRIBE v2 (d’Ascoli et al., 2026), a multimodal brain encoder selected for its fit to cortical responses across the audio, video, and text modalities; for video we used DINOv2-large, an image model applied to single frames, in place of TRIBE v2’s video backbone, V-JEPA, which encodes a window of frames at once and would carry information from neighbouring frames into each TR. For audio we used Wav2Vec-BERT 2.0 (Barrault et al., 2023); for video, DINOv2-large (Oquab et al., 2023); for text, LLaMA 3.2 3B (Grattafiori et al., 2024). All three backbones provide more than twenty layers, enough resolution for the layer-index-as-depth analyses described below (cf. the NLP-probing literature on shallow / middle / deep representations, Tenney et al., 2019; Rogers et al., 2020). We extracted features from the frozen pretrained backbones and fit no brain-prediction model to them. The decoding asks whether features learned without brain data carry information about which state is expressed; a model fit to predict these participants’ responses, even within held-out folds, would instead measure how well it predicts their activity, which state identity partly reflects by construction.

Each run of each stimulus passed through the appropriate backbone(s) once, and hidden states from every layer were saved before any downstream analysis. For audio (*Friends, Movie10, Le Petit Prince*), Wav2Vec-BERT 2.0 receives 16 kHz waveform input; per-layer activations within each 1.49 s TR window were mean-pooled into a single per-TR vector per layer. For *Le Petit Prince*, the 4 s of silence before each section were represented by two empty TRs (2.98 s), so each audio vector was assigned to a TR that began about 1 s before the sound it summarizes, and the effective feature-to-state lag for this stimulus is about 1 s shorter than the nominal lag. For video (*Friends*, *Movie10*), DINOv2-large processes a single frame sampled at each TR center, with per-layer patch-token activations mean-pooled across the spatial grid into a per-TR vector. For text (all five stimulus × language combinations), LLaMA 3.2 3B receives the cumulative transcript through TR *t* (the 128 K-token context window accommodates every run in a single forward pass); the per-TR readout uses a bounded pooling window, described below. Layers are numbered from 0, the output of the first transformer block (the embedding output is not used), to n − 1, the output of the last block, with n = 24 for Wav2Vec-BERT 2.0 and DINOv2-large and n = 28 for LLaMA 3.2 3B; relative depth is the layer number divided by n − 1, running from 0 for the first block to 1 for the last.

For text, naive pooling over the cumulative context drifts toward a running mean: each TR contributes a small fraction of new content while earlier tokens dominate. To preserve per-TR specificity, we mean-pool only over tokens whose scan-time falls inside a bounded window of *W* TRs ending at TR *t* (the forward pass still spans the full cumulative context, so each token retains access to its self-attention history; only the readout window is restricted). We chose W in two steps. First, for W ∈ {1, 3, 6, 9} TRs we checked that the pooled features still changed from TR to TR (three Friends runs, layer 14): the mean cosine similarity between features at adjacent TRs had to stay below 0.95 and exceed that between TRs paired at random within the run by at least 0.10. W = 6 and W = 9 failed (adjacent-TR similarity 0.977 and 0.988), because longer windows smooth the features toward a constant within each run; W = 1 (0.608) and W = 3 (0.925) passed. Second, in two participants (sub-01 and sub-04), with the feature-to-state lag (the temporal offset between transformer features and brain states, which absorbs the hemodynamic delay) held at 4 TRs (about 6 s, near the hemodynamic peak), we took each participant’s best layer in the layer-wise state decoding described below and measured how far its balanced accuracy exceeded the timing-regressor baseline. W = 1 gave the larger margin in both participants (smaller of the two margins 0.046 for W = 1, 0.041 for W = 3). This single fixed lag served only to select W; the decoding analysis below sweeps the lag across 0–8 TRs, and the text decoding reported in **Results** peaks at 3 TRs in every participant. We adopted *W* = 1 (single-TR mean pooling) for production extraction across all subject × stimulus cells.

Each feature dimension was first z-scored within each run, to bring feature scales into a common range across runs. For each layer we then fit a PCA on the training-split runs (the 70/15/15 season-stratified split used for fMRI dimensionality reduction), keeping the components needed to reach 95% of the explained variance, and projected the validation and test runs through the same PCA. We did not convolve transformer features with a canonical hemodynamic response; lag between transformer features and brain states is tested directly in the correspondence analyses, which sweep lag ∈ {0, 1,…, 8} TR (0–∼12 s).

To locate the model depth at which stimulus features best predicted brain-state identity, we decoded each participant’s brain-state sequence from the layer-wise transformer features (extracted above) one layer at a time. For each (participant, modality) we fit a regularized linear (ridge) multi-class classifier under leave-one-run-out cross-validation with balanced class weights, predicting the Viterbi-decoded state at each TR from that layer’s per-TR feature vector, and scored it by balanced accuracy with a normalized effect size (observed − chance) / (1 − chance). Decoding was restricted to the content-eligible state set (see **State classification**). Because the transformer layer index orders the model’s representational hierarchy from shallow to deep, the peak-decoding layer indexes the model depth whose features are most informative about brain-state identity for that modality. This is an empirical decoding readout, not a claim about where or how the brain represents that content. We report the full layer × lag grid, but lag 0, while retained as a stimulus–state synchrony diagnostic, is excluded from the peak-lag search, so every reported peak layer and lag is selected from lags 1–8.

Significance was assessed against a circular-shift permutation null: the decoded state sequence was circularly shifted within each run (1,000 permutations, the same permuted sequences reused across layers), preserving within-run temporal autocorrelation while breaking the feature–state correspondence. We applied Benjamini–Hochberg correction across the full layer × lag grid within each participant. A confound baseline reports the accuracy achievable from six timing and session-position regressors alone (a run-onset indicator, onset-timing linear, quadratic, and cubic terms, and episode and season position). We also attempted a second control that reused the run-onset-anchored states themselves as decoding targets, but it is not interpretable and we do not rely on it: those states number 2-8 against the 15-31 content-eligible targets, and normalized effect size is not comparable across different class counts, so the control scores higher than the main analysis for arithmetic rather than substantive reasons. Its label set is also heterogeneous and partly contaminated by the stereotyped series theme music. We report it for disclosure only (**Figure S6**). All inference is per participant; the across-participant medians and means shown in **Figure 5D–G** are descriptive.

To ask whether the late-layer peak of video decoding depends on particular brain networks, we repeated the within-Friends video decoding within groups of states. Each content-eligible state was assigned to the network, among the 13 functional groupings, with the largest mean absolute value in its parcel-space mean map, and labelled positive or negative by the sign of its mean over that network’s parcels, giving up to 26 groups per participant. For every group with at least five states and at least 200 TRs, we decoded state identity among the group’s states from the DINOv2-large features of each layer at the participant’s best video lag from the main analysis (3 TRs in every participant), using the same leave-one-run-out ridge classifier, a circular-shift null of 1,000 permutations generated within the group’s TRs, and Benjamini–Hochberg correction across the 24 layers within each group. Twelve groups in five participants met these criteria; none of sub-06’s groups contained five states (**Figure S5**).

To test whether this modality-specific decoding depth is preserved for other stimuli, we trained decoders on *Friends* and applied them, without retraining, to *Movie10*, *Harry Potter*, and the French and English *Le Petit Prince* runs, each treated as a separate test set. The out-of-stimulus brain activity had already been assigned *Friends* state labels by the frozen PCA + *Friends*-HMM pipeline (see **Cross-stimulus decoding**), so training and test labels come from the same state set. Test-set features were z-scored within run and projected through the per-layer PCA fitted on the *Friends* training split. For each participant, model and test set, decoding was restricted to content-eligible states with a fractional occupancy of at least 1% in both *Friends* and the test set, each computed over all of that stimulus’s TRs. At each layer, one ridge classifier (α = 1, balanced class weights) was fitted on every *Friends* TR assigned to these states, across all *Friends* runs, and applied to the test-set TRs assigned to the same states. Training and test features were taken at the same feature-to-state lag: the lag at which the participant’s within-*Friends* decoder for that model peaked (selected from lags 1–8 as above; 3 TRs for video and text in every participant, 3 or 4 TRs for audio), so the lag was chosen on *Friends* alone and never on the test set. A modality guard excluded stimulus–model pairs without the model’s input (the audio and video models for *Harry Potter*, the video model for *Le Petit Prince*). We scored balanced accuracy over the test TRs, with chance equal to one over the number of decoded states, together with the normalized effect size and per-state recall and precision. To assess significance, we circularly shifted the *Friends* training labels within each *Friends* run and refit the classifier for each of 1,000 permutations (the same shifted label sequences at every layer), evaluating each refit against the real test-set labels; this null asks whether the *Friends*-trained decoder predicts the out-of-stimulus state labels above chance. One-sided *p* values were corrected across layers within each participant, model and test set (Benjamini–Hochberg). For *Movie10*, the same decoders and permutations were also scored within each film, with correction across layers within each film.

### Use of generative AI tools

We used Claude Opus 5.5 (Anthropic) to write and revise Python code for the analysis and figure pipeline and to review manuscript drafts, including checking reported values against the analysis outputs and suggesting edits for clarity. The authors reviewed, tested and edited all code and text produced with the tool and took full responsibility for the content of the manuscript.

## Data and Code Availability

The data is available at https://www.cneuromod.ca/. The code for our analysis and visualization are available at https://github.com/yibeichan/brain-states-friends. Supplementary materials are available at https://github.com/yibeichan/brain-states-friends/tree/main/docs/supplementary

## Acknowledgement

The Courtois project on neural modelling was made possible by a generous donation from the Courtois foundation, administered by the Fondation Institut Gériatrie Montréal at CIUSSS du Centre-Sud-de-l’île-de-Montréal and University of Montreal. The Courtois NeuroMod team is based at Centre de Recherche de l’Institut Universitaire de Gériatrie de Montréal, with several other institutions involved. See the CNeuroMod documentation for an up-to-date list of contributors (https://docs.cneuromod.ca).

## Author Contributions

YC: Conceptualization, Methodology, Software, Formal analysis, Investigation, Data Curation, Visualization, Writing - Original Draft. MG: Methodology, Writing - Review & Editing. MSL: Resources, Data Curation, Writing - Review & Editing. LB: Resources, Writing - Review & Editing. SG: Conceptualization, Funding acquisition, Methodology, Resources, Writing - Review & Editing, Supervision.

